# Fear of grazing rivals the toxin induction effect of nitrogen enrichment in marine harmful algae— a meta-analysis

**DOI:** 10.1101/2025.03.07.642069

**Authors:** Milad Pourdanandeh, Erik Selander

## Abstract

One of the major subfields of chemical ecology is the study of toxins and how they mediate interactions between organisms. Toxins produced by harmful algae, phycotoxins, impact a wide variety of organisms connected to the marine food web. Significant research efforts have thus aimed to identify the ecological and evolutionary drivers behind harmful algal blooms (HABs) to facilitate their forecasting, mitigation, and management. Nutrient availability is a key factor controlling growth and toxin production. Additionally, recent evidence has shown that harmful algae can sense the presence of zooplankton grazers, primarily copepods, and respond by dramatically increasing toxin production. Phycotoxin production is consequently controlled by a combination of bottom-up and top-down drivers, but the relative importance of the two is not understood. Here we conducted a meta-analysis of 113 control-treatment contrasts from 37 peer-reviewed experimental studies, comparing the effects of relative nitrogen enrichment (increased nitrogen:phosphorus ratio) and elevated grazing risk on phycotoxin induction in the two most studied marine HAB-forming genera, *Alexandrium* dinoflagellates and *Pseudo-nitzschia* diatoms. We show that phycotoxins are induced in response to both nitrogen enrichment and elevated grazing risk. Although both genera responded similarly to nitrogen enrichment, *Pseudo-nitzschi*a toxins increased four times more in response to grazers than to nitrogen enrichment, and ten times more than *Alexandrium* toxins did in response to grazers. Grazing risk thus appears to rival, perhaps even supersede, the well-established phycotoxin-inducing effect of nitrogen enrichment in marine harmful algae. Although this analysis is limited to the two most studied marine HAB genera, we conclude that future attempts to understand the evolution and variable production of phycotoxins require integration of bottom-up nutrient availability and top-down selective pressures to fully elucidate phycotoxin dynamics in marine HAB-forming species.

## 1. Introduction

Algal blooms are key features of oceanic primary production, contributing to significant bursts in production (Anderson, 1989; Smayda, 1989). Of the 5000 known species of extant marine phytoplankton, less than 10% bloom to such extent as to notably discolour the sea surface and approximately 2–4% produce toxins (phycotoxins) that can harm humans (Hallegraeff, 1993; Landsberg, 2002). Further, the number of marine phycotoxin producers has increased with improved identification technologies and global monitoring efforts in recent decades (Landsberg, Van Dolah & Doucette, 2005; Hallegraeff *et al*., 2021).

Consequently, significant research efforts have been invested into understanding the factors that control the abundance of harmful algae and their highly variable toxin production capacity. Phycotoxins encompass a rich diversity of compounds and the evolutionary rationale for their production is still debated (Cembella, 2003; Pančić & Kiørboe, 2018).

Eutrophication, particularly nitrogen enrichment, is widely recognized as a key driver of increased cellular phycotoxin content in harmful algae (Boyer *et al*., 1987; Hallegraeff, 1993; Brandenburg *et al*., 2020). However, mounting evidence suggests that some phycotoxins— mainly alkaloids such as domoic acid and saxitoxins—are induced by zooplankton grazers and serve as chemical defences against them (Pančić & Kiørboe, 2018).

### 1.1 The raison d’être of plant secondary metabolites

Although morphological defences, such as spines and thorns, have historically been recognised as adaptations evolved to deter herbivores (Karban & Baldwin, 1997), ecologists in the early 1900s were more hesitant to acknowledge grazer-deterring functions of compounds not involved in primary (i.e. secondary metabolites, Fraenkel, 1959; Hartmann, 2008). This perspective persists to this day, partly due to the consistent framing of these compounds as ‘secondary’ to the primary metabolites involved in development, growth, and reproduction of organisms (Tversky & Kahneman, 1981; Matthes & Schemer, 2012). Several theoretical frameworks have been developed to explain and predict why many plants, and other photosynthetic eukaryotes, produce and allocate resources to secondary metabolites. These can be divided into resource-driven and demand-driven models (Stamp, 2003), which both emerged concurrently in the early 1980s. However, resource-based models—notably the influential *carbon:nutrient balance* model—found wider immediate support across most study systems (Karban & Baldwin, 1997).

The *carbon:nutrient balance* model states that the production of secondary metabolites, including potential defence chemicals, is governed by the relative availability of carbon and nutrients (Bryant, Chapin & Klein, 1983; Bryant *et al*., 1989; Reichardt *et al*., 1991) and assumes that primary growth processes take precedence over secondary defence functions (Karban & Baldwin, 1997; Koricheva *et al*., 1998). When resource availability exceeds the requirements of primary metabolism, plants can divert the surplus to the production of secondary defence metabolites (Herms & Mattson, 1992). For example, under nutrient-limited conditions, plants may allocate excess carbon to carbon-based defences, such as polyphenols (Bryant *et al*., 1989) at a neglectable cost as growth is limited by the lack of other elements regardless. The core predictions of the *carbon:nutrient balance* model is supported by evidence that carbon gain and growth depend on a plant’s mineral nutrient reserves (reviewed by Tuomi and colleagues (1988)) and that growth is more inhibited by nutrient limitation than photosynthesis (reviewed by Luxmoore (1991) and Herms & Mattson (1992)). It also acknowledges that carbon surplus can be allocated to defence (Tuomi *et al*., 1988). Over 200 studies have examined the *carbon:nutrient balance model*, a meta-analysis of 147 of these suggests that pooled non-nitrogen containing secondary metabolites and carbohydrates respond to shade, nutrient, and CO_2_ variation as predicted (Koricheva *et al*., 1998). Importantly however, many of the primary studies are riddled with methodological issues (detailed by Stamp (2003)) which may result in misleading syntheses. In summary, resource-based models have provided fundamental mechanistic insights into plant secondary metabolite production.

Demand-based models such as the *optimal defence* model (Rhoades, 1979; Roff, 1992; Stearns, 1992) builds on the terrestrial plant-herbivore interaction experiments pioneered by Ernst Stahl (1888) and entomologists in the 1950s (Fraenkel, 1959; Hartmann, 2008). While these models were primarily developed to explain secondary metabolites in terrestrial systems, they later found support in macroalgae-herbivore systems as well (Cronin & Hay, 1996; Pavia *et al*., 1997; Pavia & Toth, 2000; Rohde, Molis & Wahl, 2004). In contrast to resource-based models, demand-based models rely on an optimality approach in relation to grazing pressure. The *optimal defence* model focuses on costs and benefits of defence expressions *per se*, emphasising how natural selection shapes the adaptive allocation of defences in response to herbivory (Rhoades, 1979; Zangerl & Bazzaz, 1992). A key assumption of the *optimal defence* model, shared with other demand-based models, is that defences are costly and produced at the expense of growth and reproduction (Rhoades, 1979; Coley, Bryant & Chapin, 1985a; Simms & Rausher, 1987). Plants should therefore produce defence chemicals in proportion to the risk and severity of herbivore attacks, while balancing these needs against the metabolic costs of producing the defences (Fagerström, Larsson & Tenow, 1987; Karban, Agrawal & Mangel, 1997). In addition to explaining the evolution of defence compounds, it also elucidates the allocation of these limited resources on ecological time scales. A key strategy predicted by the *optimal defence* model is inducible defence expression, where plants increase defences only in response to grazing or reliable cues of increased grazing risk, thereby conserving resources when threats are absent (Karban & Myers, 1989; Tallamy & Raupp, 1991). High-risk or high-value plant parts are thus expected to receive greater protection, and allocation to defence is minimised when herbivore pressure is low. Demand-based models thus aim to describe how plants evolve and optimally allocate costly defences in response to herbivory risk, providing an evolutionary framework that captures the fine-scale temporal dynamics of plant-grazer interactions. The *optimal defence* model is supported by evidence of high intraspecific genetic variation in secondary metabolite type and amounts (Dirzo & Harper, 1982; Zangerl & Berenbaum, 1990), and by evidence that herbivores act as strong selective pressures for plant traits that reduce herbivory (Simms & Rausher, 1989; Mauricio & Rausher, 1997). It is also supported by evidence that defences are allocated proportionally to the risk of herbivory (Baldwin & Karb, 1995; Zangerl & Rutledge, 1996), that defences incur allocation costs (Vrieling & van Wijk, 1994; Strauss *et al*., 2002), and that induced defences enhance plant fitness (Baldwin, Sims & Kean, 1990; Agrawal, 1998).

In addition to the more influential *carbon:nutrient balance* and the *optimal defence* models described above, the *growth rate* (Coley, Bryant & Chapin, 1985b; Coley, 1988), the *plant apparency* (Feeny, 1976), and the expanded *growth-differentiation balance* (Loomis, 1932; Herms & Mattson, 1992) hypotheses are also empirically supported to varying degrees and have been valuable frameworks for organizing and directing research on plant defences. Consequently, the field of plant defences stands out in ecology for its wealth of coexisting hypotheses that have been developed and tested over several decades, without any being conclusively rejected. Berenbaum (1995) lists more than ten different hypotheses to account for the allocation of chemical defences in plants, which was further added to by Herms and Mattson (1992). For a comprehensive overview and comparison of plant defence hypotheses/models, see Stamp (2003).

### 1.2 Evidence from the pelagic: zooplankton-phytoplankton interactions

Research on toxins of harmful algae in pelagic aquatic systems has in many ways paralleled that of terrestrial plants, initially focusing more on resource-based mechanisms. This has revealed consistent relationships between nutrient dynamics and phycotoxin production (Boyer *et al*., 1987; Smayda, 1997; Anderson, Glibert & Burkholder, 2002; Glibert, 2017). Specifically, that the relative availability of nitrogen (N) and phosphorus (P) affects the amounts of phycotoxins produced in a predictable manner (Brandenburg *et al*., 2020). Higher relative nitrogen abundance typically increases the formation of nitrogen-rich phycotoxins such as domoic acid and paralytic shellfish toxins, as well as cyclic peptides such as microcystin. This aligns well with the *carbon:nutrient balance* model in showing how limiting nutrients can favour or inhibit the production of secondary metabolites. The toxin-inducing effects of nitrogen has especially contributed to the dominance of eutrophication as an explanatory mechanism for HAB-formation and expansion (Anderson *et al*., 2002; Hallegraeff, 2003; Wells *et al*., 2020). The evolutionary drivers favouring phycotoxin production and dynamics, while harder to identify under many resource-based models, are frequently discussed in the HAB literature (Cembella, 2003; Granéli & Turner, 2006; Pohnert, Steinke & Tollrian, 2007). Similarly, the production of noxious secondary metabolites has been debated from both physiological and evolutionary perspectives. From a physiological standpoint, potential defence chemicals have historically been viewed as little more than metabolic waste products (Robinson, 1974; Haslam, 1985; Waterman & Mole, 2019), although their potential defensive properties were never explicitly excluded. While resource-based factors (particularly nitrogen and phosphorus availability) have been central to explanations of phycotoxin production, increasing evidence of an evolutionary “chemical arms race” (Dawkins & Krebs, 1979; Stearns, 1992; Karban *et al*., 1997; Smetacek, 2001) with grazing zooplankton highlights the important role of grazers in further shaping their production dynamics.

Studies on zooplankton-mediated defence traits in phytoplankton have increased in recent decades. The idea that secondary metabolites may protect algae from zooplankton grazers is not new (Sykes & Huntley, 1987). More recent experiments, however, provided new empirical support for this by demonstrating that some phytoplankton species can detect copepods—the most abundant mesozooplankton grazers in marine environments—and respond by expressing a variety of physiological, morphological, and behavioural defences (Guisande *et al*., 2002; Selander *et al*., 2006, 2011; Pondaven *et al*., 2007; Pančić & Kiørboe, 2018). Copepod-induced phycotoxin production was first demonstrated in co-culture experiments, showing that direct exposure to grazers can stimulate paralytic shellfish toxins in harmful dinoflagellates such as *Alexandrium* spp. (Guisande *et al*., 2002; Selander *et al*., 2006; Bergkvist, Selander & Pavia, 2008; Senft-Batoh *et al*., 2015) and amnesic shellfish toxins in diatoms such as *Pseudo-nitzschia* spp. (Tammilehto *et al*., 2015; Harðardóttir *et al*., 2015; Lundholm *et al*., 2018). In addition to these direct effects of copepod exposure, harmful algae respond similarly to chemical cues exuded by copepods, specifically a group of taurine-conjugated polar lipids known as copepodamides (Selander *et al*., 2015; Grebner *et al*., 2019). Copepodamides appear to be ubiquitous in both marine and limnic copepods (Arnoldt *et al*., 2024) and serve as a general alarm cue for their prey in marine ecosystems (Wohlrab, Selander & John, 2017; Lindström *et al*., 2017; Selander *et al*., 2019; Arias *et al*., 2021; Rigby *et al*., 2022). However, some HAB-formers, such as *Dinophysis*, appear less responsive to copepodamides (Pourdanandeh *et al*., 2025). In response, copepods can discriminate against better-defended cells (Huntley *et al*., 1986; Teegarden, 1999; Olesen *et al*., 2020; Ryderheim, Selander & Kiørboe, 2021), providing a direct fitness benefit for algae that tailor their toxin production according to grazing risk. This selective avoidance demonstrates the evolutionary advantage of inducible defences, as increased algal toxicity reliably reduces grazing by copepods (Selander *et al*., 2006; Wohlrab *et al*., 2017; Ryderheim *et al*., 2021; Olesen *et al*., 2022). The few comparative studies of both grazer-mediated and nutrient-mediated effects on dinoflagellate paralytic shellfish toxin production suggest that, while nitrogen and phosphorus availability affects toxin production, grazer-induced effects prevail in a broad range of nutrient conditions (Selander, Cervin & Pavia, 2008; Griffin, Park & Dam, 2019). However, the relative importance of nitrogen enrichment and grazing on phycotoxins is still not known.

### 1.3 Rationale and aim

The recent surge in grazer-mediated phycotoxin induction studies has now reached a critical mass, enabling synthesis and direct quantitative comparisons to the well-established induction effects of nitrogen enrichment. The aim of this meta-analysis was therefore to quantify and compare the effects of nitrogen enrichment and grazing risk on phycotoxin production in HAB-forming marine algae. We targeted two genera, *Alexandrium* dinoflagellates and *Pseudo-nitzschia* diatoms (Fig. 1) as they represent the most extensively studied genera in respect to both resource and demand driven processes. Nitrogen enrichment (resource) was defined as an increased nitrogen:phosphorus (N:P) ratio, and grazing risk (demand) broadly as direct or indirect exposure to algae-grazing zooplankton or their chemical cues.

**Fig. 1.**
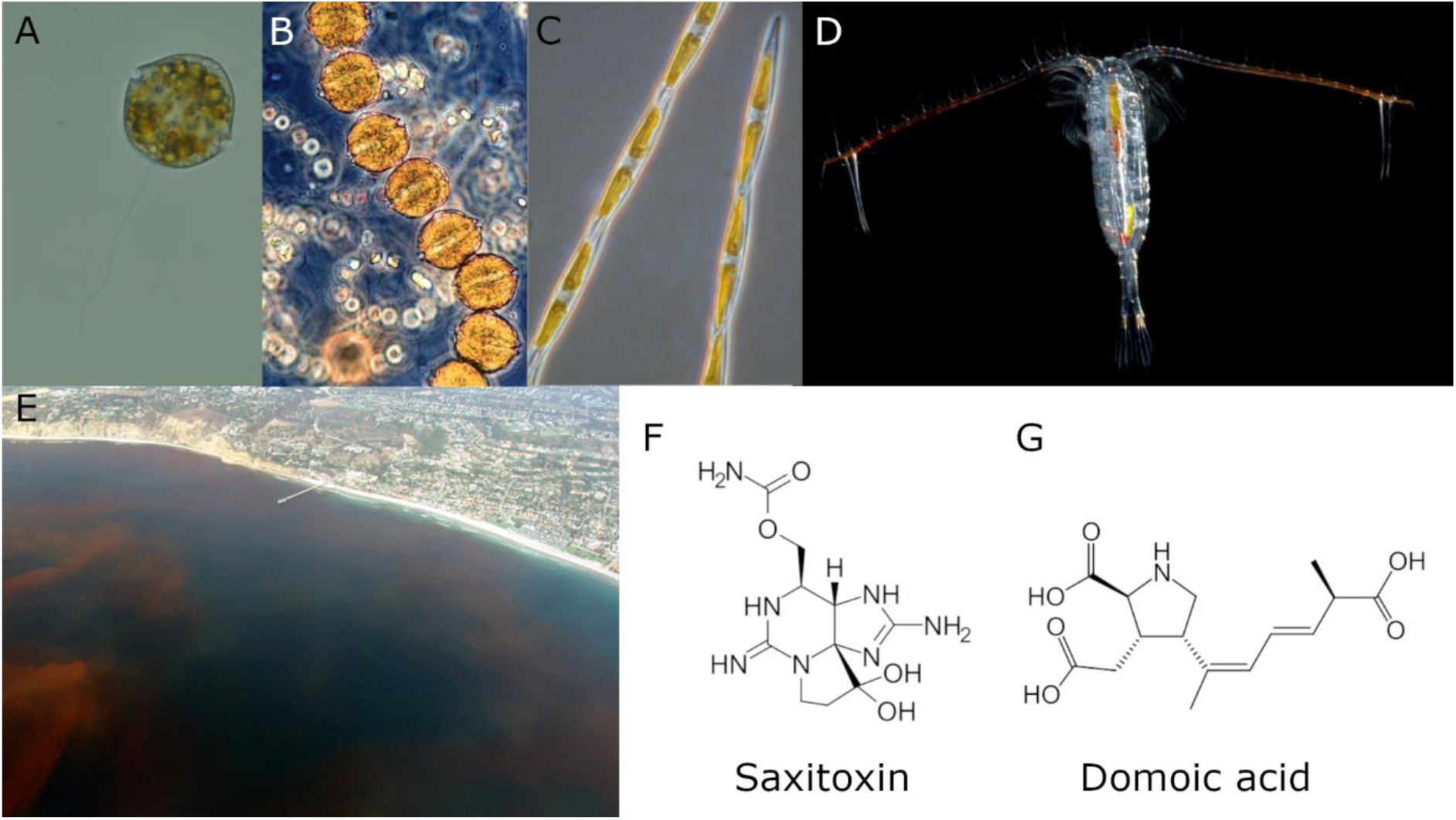
The primary organisms, and phycotoxins examined in this meta-analysis. (A) Single cell of the dinoflagellate *Alexandrium catenella*. (B) Chain of *A. catenella* cells. (C) The pennate diatom *Pseudo-nitzschia* sp. (D) A *Calanus* sp. copepod, a common marine mesozooplankton grazer in the Atlantic. (E) A dinoflagellate-driven red tide off the Scripps Institution of Oceanography Pier, La Jolla, California. (F) Chemical structure of saxitoxin, one of the principal paralytic shellfish toxins produced by some HAB-forming dinoflagellates. (G) Chemical structure of domoic acid, the phycotoxin produced by *Pseudo-nitzschia* that causes amnesic shellfish poisoning (ASP) syndrome. Photo credits: (A) Fredrik Ryderheim, (B) Jan Rines, (C) Rozalind Jester, (D) Erik Selander.

## 2. Materials & methods

### 2.1 Literature search and screening

We used Web of Science (Clarivate Analytics, 2021-03-18) to search for relevant studies using two search strings. For studies on toxin induction in response to grazers or grazer chemical cues, designated as “demand” or “grazing” papers and effects, we used “*induc* AND toxi* AND ((alexandrium OR pseudo-nitzschia) AND (graz* OR copepod* OR zooplank*)*” as search terms. For studies on toxin induction as a response to changes in nitrogen and/or phosphorous load, designated “resource” or “nutrient” papers and effects, we used “*induc* AND toxi* AND (nitr* OR phosph*) AND (alexandrium OR pseudo-nitzschia) AND (produc* OR synthes*)*”. The searches yielded 281 papers, 48 demand and 233 resource, for screening. Additionally, 14 potentially relevant papers were added to the initial pool of records; 12 relevant papers used by Brandenburg and colleagues (2020) in their meta-analysis, and two known papers from closely connected research groups that the search missed. After removing ten duplicates, this yielded a total of 285 papers for screening. Based on titles and abstracts, 95 papers were identified as relevant for full-text assessment, excluding the 188 papers that did not study toxin production in our chosen taxa, and two papers that we were unable to retrieve full-text versions of (Fig. 2).

**Fig. 2.**
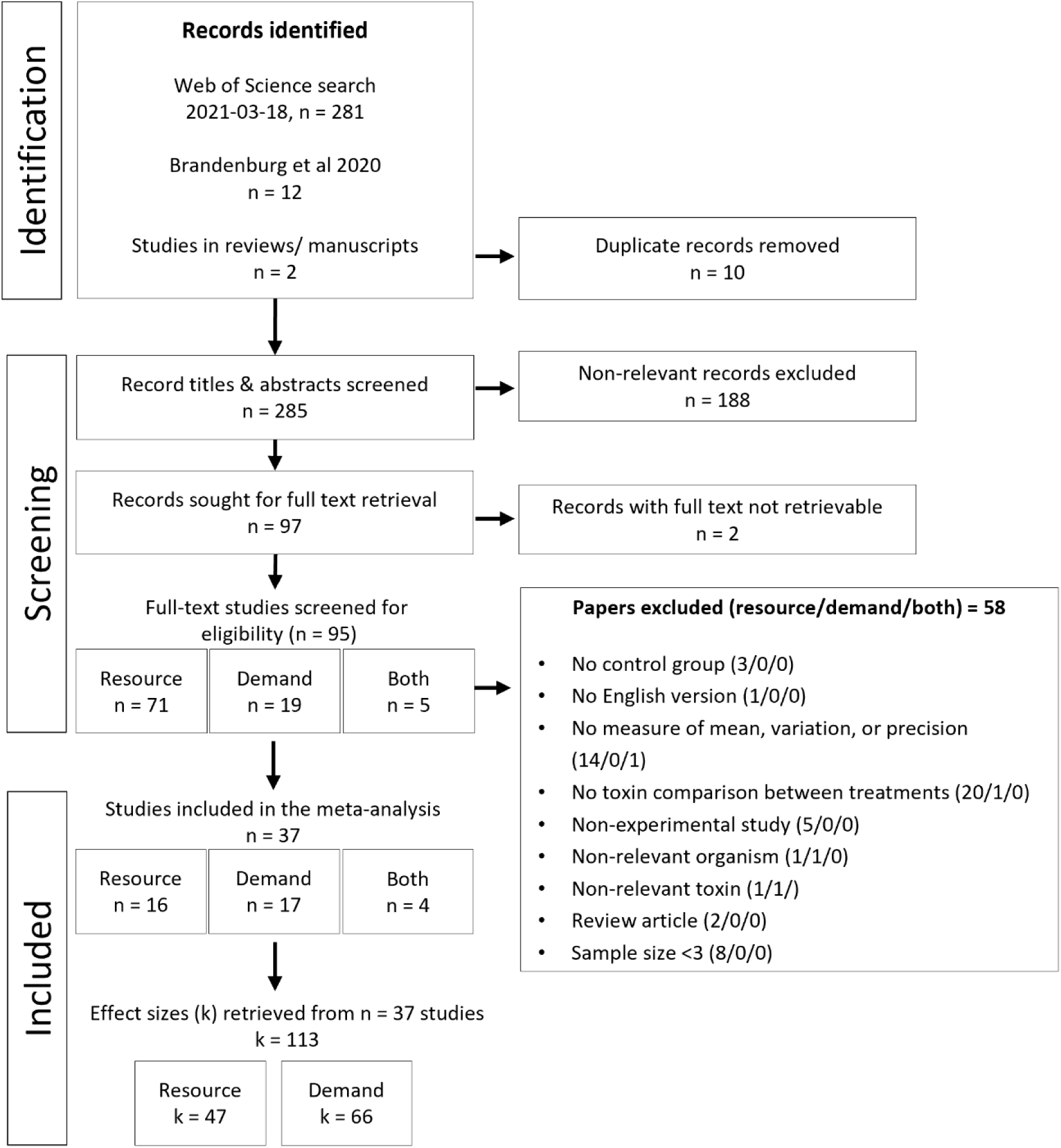
Preferred Reporting Items for Systematic Reviews and Meta-Analyses (PRISMA) flow chart showing the number of articles found, screened, and included/excluded. r = resource (nitrogen enrichment), d = demand (increased grazing risk).

### 2.2 Eligibility criteria

From the 95 full-text papers screened for eligibility, we excluded experimental studies shorter than 48 hours that did not assess the effects of zooplankton or zooplankton chemical cues or different nutrient loads (nitrogen:phosphorus ratio) on toxin production/content for *Alexandrium* or *Pseudo-nitzschia*. We also required presentation of descriptive statistics (mean, measure of variance/confidence and sample size) and a minimum sample size of three for each experimental group. If unavailable, we requested the raw or summarised data from the authors (data received for 8 studies). If the authors could not provide this, or did not respond, we extracted data from figures (n = 7 studies) with PlotDigitizer version 2.2 (2022).

In factorial experiments, data extraction was limited to ambient/control levels of the other factor (e.g., light or temperature).

Demand effects were only included if there was a control treatment without grazers or grazer cues, and a grazer treatment with zooplankton or their chemical cues added.

Additionally, if both direct and indirect grazing effects were assessed in the same experiment, e.g., if cells from both inside and outside of a cage containing grazing zooplankton were analysed for toxins, only the indirect (outside) effect was used. This was done because the indirectly affected cells should, in theory, contribute more conservatively to the overall mean effect in the meta-analysis compared to the cells grazed on directly. Studies on resource effects were included only if they contained a minimum of two independent experimental treatments with different nutrient loads. Excluding the papers that violated these criteria resulted in 37 papers used for analyses.

### 2.3 Data extraction and effect size calculation

From the 37 papers identified, we extracted means, standard deviations, and sample sizes from each paper, resulting in 113 control-treatment contrasts. If precision estimates (standard error or confidence intervals) were presented instead, these were extracted and used to calculate standard deviations. We also extracted moderator (predictor) information/metadata, e.g., culturing temperature, salinity, light regime, difference in N:P ratio etc., to assess underlying effects of experimental conditions on our synthesised effects. If experiments included multiple treatment groups, such as varying levels of relative nitrogen enrichment, grazer densities, or grazer cue concentrations, we extracted only the maximum effect contrast to capture the greatest induction range for each driver level (resource and demand). For resource effects, the treatment with the lower N:P ratio was consistently used as the control, the treatment group thus represented nitrogen enrichment effects relative to the control. We calculated and used the small sample bias-corrected log response ratio (LRR^Δ^) proposed by Lajeunesse (2015) as effect size. LRR^Δ^ is defined as:

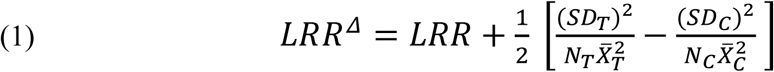

where 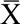, SD, and N are the means, standard deviations, and sample sizes of the treatment (T) and control (C) groups, respectively. LRR is the standard log response ratio (Hedges, Gurevitch & Curtis, 1999) given by equation:

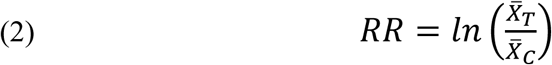

where 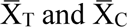 are the mean values of the treatment and control groups, respectively. The sampling variance of LRR^Δ^, var(LRR^Δ^), is given by equation:

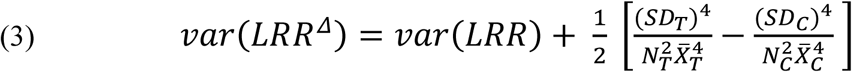

where var(LRR) is defined as:

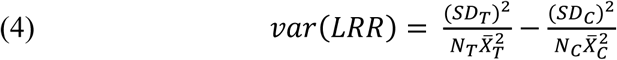

### 2.4 Meta-analyses and meta-regressions

We used multi-level mixed effect meta-analytical models to evaluate the effects of grazing risk (demand) and relative nitrogen enrichment (increased N:P or resource) on phycotoxin production in *Alexandrium* and *Pseudo-nitzschia*. These were implemented in R using the *metafor* package (Viechtbauer, 2010) and constructed to treat effects within each driver (demand or resource) as random but effects between drivers as fixed. This reflects the fact that demand (top-down induction) and resource (bottom-up induction) represent the only two possible experimental categories of interest in this analysis, unlike moderators such as species, which could encompass levels beyond those studied here. This enabled us to estimate and account for heteroscedastic variances among moderator levels, avoiding the common assumption of random-effects models that all moderator levels share equal variances, which is rarely satisfied in ecological studies (Senior *et al*., 2016). To account for hierarchical structure and variability, random effects were included for study ID, capturing between-study variance, and effect ID, modelling residual heterogeneity within studies. I² statistics were calculated to quantify the proportion of total variance attributable to heterogeneity beyond sampling error (Higgins & Thompson, 2002). The models were constructed with code adapted from Nakagawa and colleagues (2021, 2023).

Meta-regressions were used to test the influence of continuous variable moderators such as temperature, salinity etc. Moderators were added individually to univariate models to evaluate their impact on effect size variability, while multi-moderator models assessed combination effects. These included interactions between driver (demand vs. resource) and environmental or experimental moderators to explore potential context-dependent dynamics in phycotoxin production. Detailed results of these univariate and two-way moderator interaction analyses are available in subsection 5.1 of the Supplementary code. Several models were unable to reach convergence or had other issues we were unable to solve.

Therefore, some potential interaction effects have not been identified.

### 2.5 Sensitivity analyses

Response ratios and their variances are not accurate approximations of the response ratio sampling distribution when primary study sample sizes are small (Hedges *et al*., 1999), and when the mean value of either control or treatment group is close to zero (Lajeunesse, 2015), which can bias the result outcome of the overall synthesis. We therefore used a modified version of Geary’s test (Geary, 1930; Hedges *et al*., 1999) proposed by Lajeunesse (Eq. 13: 2015) to identify these potentially problematic effect sizes. We omitted these to determine their influence on the overall results and whether they should be removed from the final analysis.

### 2.6 Publication bias

Publication bias encompasses several types of bias related to the dissemination of scientific information. Here, we focus on two main types: outcome reporting bias, which occurs when studies are selectively published (the file drawer problem; Rosenthal, 1979), and time-lag bias, where significant, large-effect, or corroborating results tend to be published earlier than non-significant, incremental, or negative findings (i.e. the decline effect: Koricheva & Kulinskaya, 2019; Connell & Leung, 2023). Two features common to meta-analyses in ecology and evolution, large amounts of heterogeneity and non-independence of effects, complicate and invalidate many standard methods for detecting and quantifying bias. Due to substantial heterogeneity and non-independent effects in our dataset—mainly driven by differential research effort across taxa, methodological variation between experiments, and effect sizes being nested within both primary studies and research groups—conventional meta-analytic diagnostic approaches such as funnel plots, trim-and-fill procedures, and leave-one-out analyses are unsuitable. Following recommendations by Nakagawa and colleagues (2022), we therefore used a cumulative multi-level meta-analysis to visually assess time-lag bias by examining changes in mean effects over time (Harvard & Lau, 1993; Lau, Schmid & Chalmers, 1995). This was complemented by multilevel meta-regressions to simultaneously test for publication bias using the square root of the inverse “effective sample size” as predictor (Nakagawa *et al*., 2022), and time-lag bias using publication year as a predictor (Jennions & Møller, 2002; Koricheva & Kulinskaya, 2019).

### 2.7 Software and statistics

All analyses and data visualisations were performed in *R* v.4.4.1 (R Core Team, 2024) using RStudio v. 2024.4.1.748 (Posit team, 2024) and packages: *orchaRd* (Nakagawa *et al*., 2021, 2023), *ggforestplot* (Scheinin *et al*., 2023), *tidyverse* (Wickham *et al*., 2019), *readr* (Wickham, Hester & Bryan, 2024), *devtools* (Wickham *et al*., 2022), *glmulti* (Calcagno, 2020), *patchwork* (Pedersen, 2024), *multcomp* (Hothorn, Bretz & Westfall, 2008), *emmeans* (Lenth, 2024), *metafor* (Viechtbauer, 2010), *ggtext* (Wilke & Wiernik, 2022), *ggridges* (Wilke, 2024). The analysis code, and the data sets it uses to perform all analyses and generate all figures, are openly accessible at https://doi.org/10.5281/zenodo.14713104 (Pourdanandeh & Selander, 2025). Significance level ɑ = 0.05 was used for all statistical tests. Estimated effects are presented as percentage increases (%) from control, with 95% confidence intervals given in parentheses, e.g. 100% (70–150). 95% prediction intervals, i.e. where 95% of new effect sizes are predicted to fall with repeated sampling of the literature, are presented in figures only. Note that confidence and prediction intervals are asymmetric around estimated means, due to being derived from a natural log-scale.

## 3. Results

Of the 113 effect sizes (k) extracted from 37 eligible studies (n) on phycotoxin induction, 66% were from experiments on *Alexandrium* and 34% on *Pseudo-nitzschia* (Supplementary Fig. S1). Resource effects (elevated N:P ratio) accounted for 42% of total effect sizes (*Alexandrium:* 26%, *Pseudo-nitzschia:* 16%), while demand effects (grazer) comprised the remaining 58% (*Alexandrium:* 41%, *Pseudo-nitzschia:* 18%; Fig. S2). The primary studies involved 35 phytoplankton strains (Fig. S3) representing 9 species (Fig. S4). *Alexandrium minutum* and *Pseudo-nitzschia seriata* comprised the majority of effect sizes within their respective genera, accounting for 49% and 53% of estimates, respectively. Experiments which exposed phytoplankton to live grazers were conducted on 13 zooplankton taxa (Fig. S5). The studies were published between 1999 and 2022, with the majority (51%) published after 2015, accounting for 62% of total effect sizes (Fig. S6). Sample sizes ranged from 3 to 20 replicates (mean = 3.5, median = 3, mode = 3) and showed no change over time (Fig. S7). Almost all studies included in the analysis used batch cultures, which accounted for 89% of effect estimates (Fig. S8). Notably, all experiments that used continuous or semi-continuous methods were conducted on *Alexandrium.* Illumination level (photon flux) in the experiments ranged from 50 to 350 µmol m⁻² s⁻¹, but the majority (68%) were conducted at levels between 80 and 150 µmol m⁻² s⁻¹ (Fig. S9A). Light:dark regimes ranged from 12 to 22 hours of light per day, with 73% of experiments using either 12 hours (50%) or 14 hours (23%) of light (Fig. S9B). Experimental temperature regimes ranged from 4 to 25 °C, with most (k = 74) conducted at 16 to 18 °C (Fig. S9C). Salinity was among the least consistently reported variables, with 36% of effects (k = 41) missing salinity data. Reported salinities ranged from 15 to 35 PSU, with the majority (44%) between 33 and 35 PSU (Fig. S9D). Experiment durations varied from 2 to 240 days, but most were shorter than 7 days, with a median duration of 5 days (Fig. S9E).

### 3.1 Main effects

Both nitrogen enrichment (resource) and elevated grazing risk (demand) significantly increased phycotoxins by 267% (95% CI: 125–498) and 388% (230–622), respectively (Fig. 3, Table 1:Ⅰ). Although elevated grazing risk increased phycotoxin levels by 121 percentage units more than nitrogen enrichment, they did not significantly differ from one another (*p* = 0.35, Table 1). An interaction effect emerged when phytoplankton genus was included as a moderator (*p* = 0.02, Table 1:ⅠⅠ), i.e. the effect of driver differed between the two genera.

**Fig. 3.**
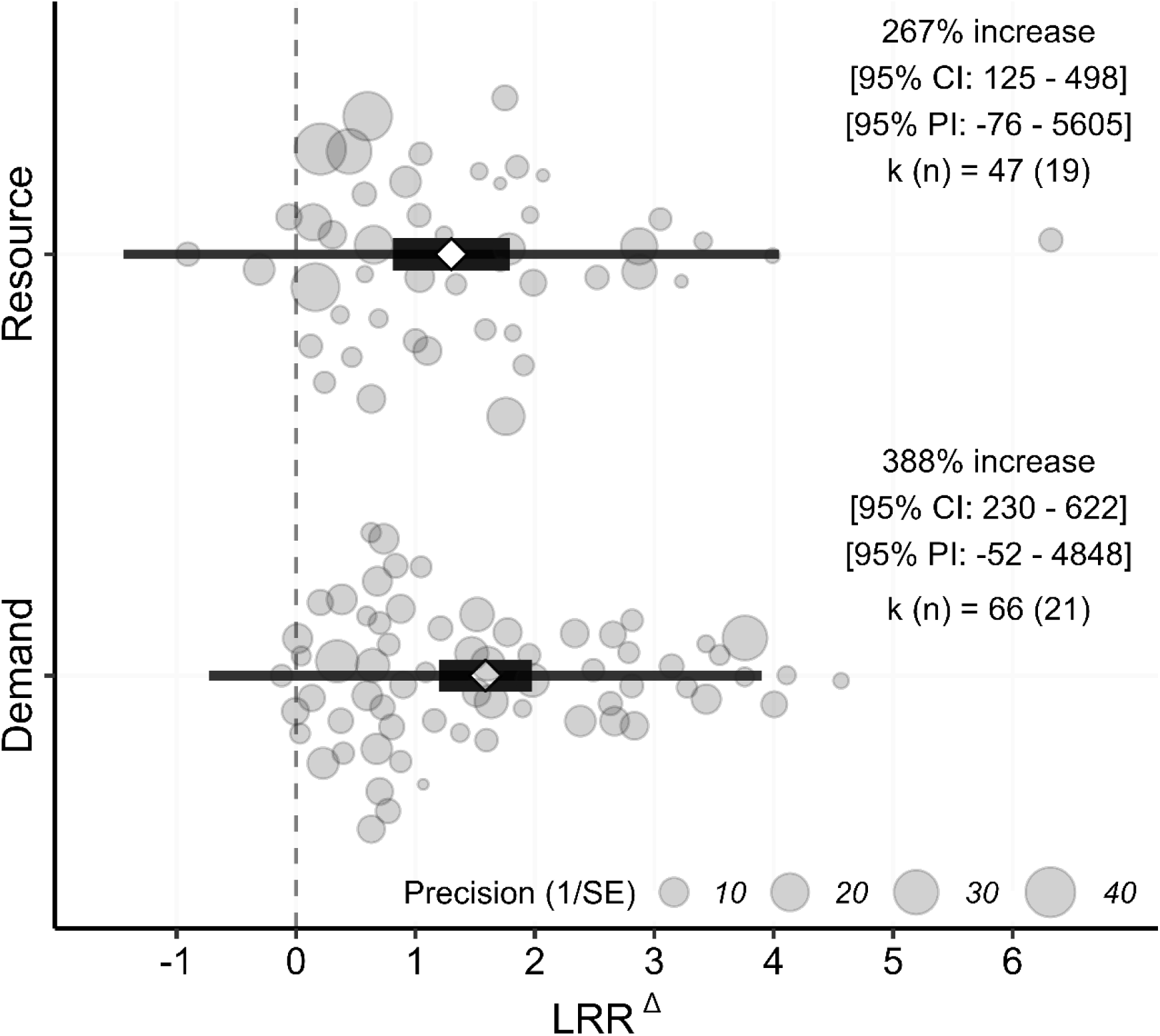
Effects of relative nitrogen enrichment (resource) and elevated grazing risk (demand) on phycotoxin induction in two harmful algal bloom-forming marine phytoplankton genera. Effect size (x-axis) is the small sample bias-corrected log response ratio (LRR^Δ^) proposed by Lajeunesse (2015). White diamonds represent the mean effect size, thick black boxes denote the 95% confidence intervals (95% CI) of the mean effect, and thin black lines denote the 95% prediction intervals (95% PI; where 95% of new effect sizes are expected to fall with repeated sampling of the literature). Circles denote individual effect sizes (k) from a given number of studies (n), sized inversely proportional to their sampling error (1/SE). Back-transformed mean values, 95% CI limits, and 95% PI limits are presented as corresponding percentages in the annotated text.

**Table 1.**
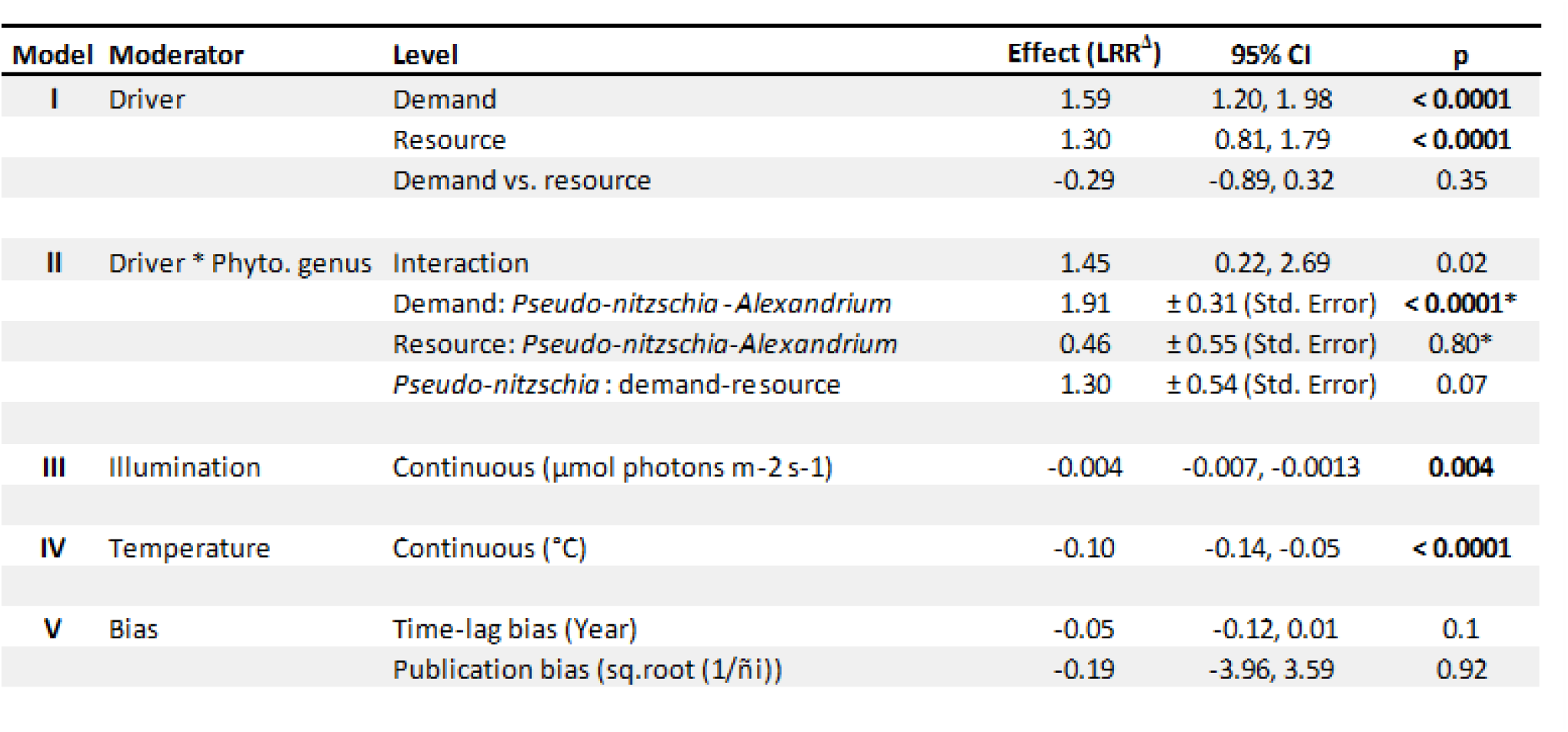
Results from meta-analytical models referenced in the main text. The models correspond to the following in the Supplementary code: I = mod_13, II = mod_16, III = mod_8, IV = mod_10, and V = bias_all. Results of pairwise level contrasts for model II are presented as estimated differences ± standard errors, with p-values adjusted (*) using Holm’s sequential multiple test procedure (Holm, 1979). Effects are expressed as small-sample bias-corrected log response ratios (LRR^Δ^), as proposed by Lajeunesse (2015). Statistically significant model terms (p ≤ 0.05) are highlighted in bold. The term ñi denotes the “effective sample size” described by Nakagawa and colleagues (2022) and used as a predictor (moderator) when testing for publication bias.

Phycotoxin induction due to increased grazing risk was 10-fold higher in *Pseudo-nitzschia* compared to *Alexandrium* (*p* < 0.0001, Table 1, Fig. 4) while both genera were similarly affected by nitrogen enrichment (*p* = 0.8, Fig. 4). In addition, demand effects on *Pseudo-nitzschia* toxins were 4-fold higher than resource effects, but this difference was marginally non-significant (*p* = 0.07, Table 1).

**Fig. 4.**
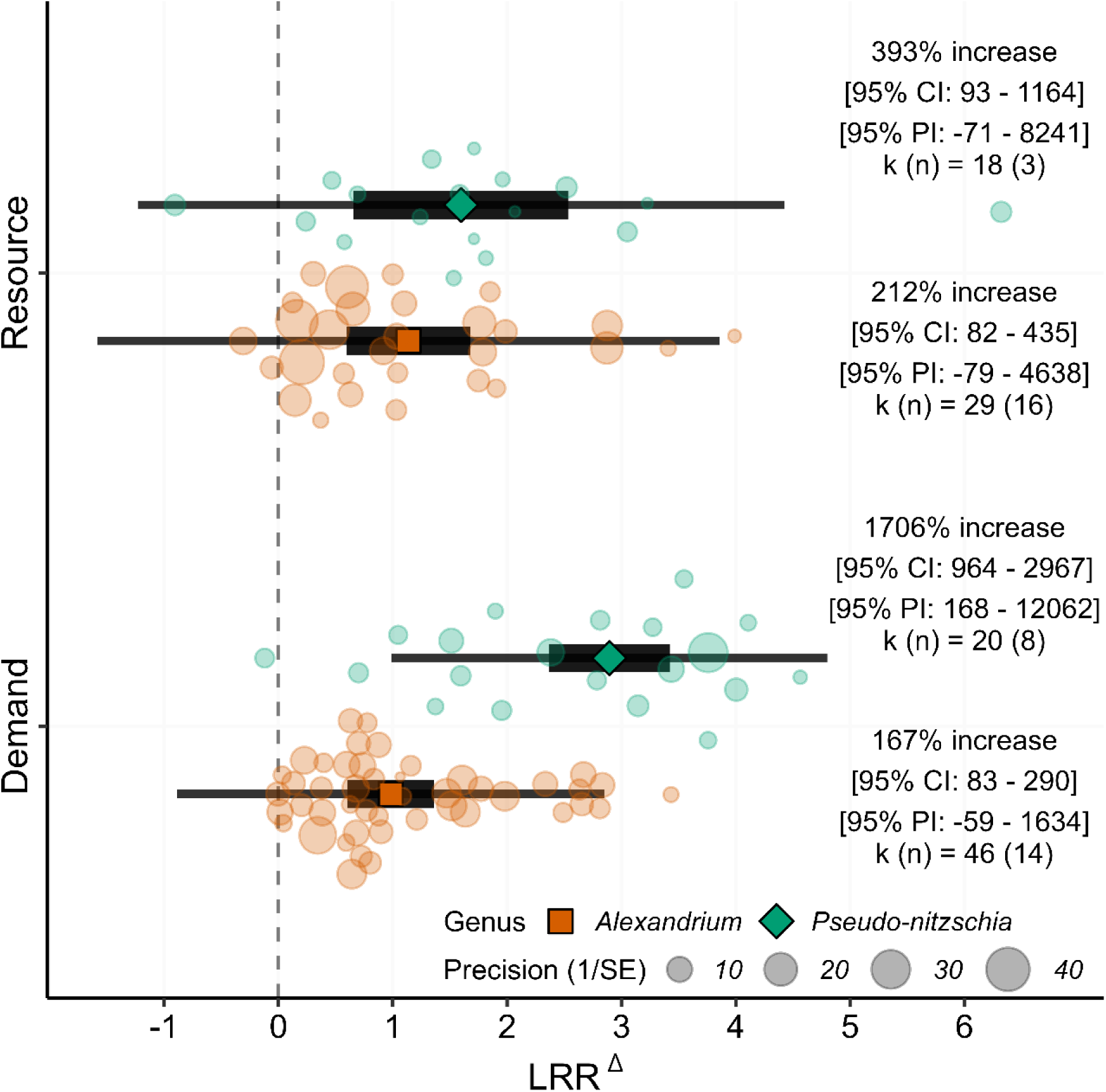
Effects of relative nitrogen enrichment (resource) and elevated grazing risk (demand) on phycotoxin induction, partitioned by phytoplankton genus (*Alexandrium*: orange; *Pseudo-nitzschia*: green). Effect size (x-axis) is the small sample bias-corrected log response ratio (LRR^Δ^) proposed by Lajeunesse (2015). Orange squares represent mean effect sizes for *Alexandrium*, and green diamonds represent mean effect sizes for *Pseudo-nitzschia*. Thick black lines denote 95% confidence intervals (95% CI) for the mean effects, and thin black lines denote 95% prediction intervals (95% PI; where 95% of new effect sizes are expected to fall with repeated sampling of the literature). Circles denote individual effect sizes (k) from a given number of studies (n), sized inversely proportional to their sampling error (1/SE). Back-transformed mean values, 95% CI limits, and 95% PI limits are presented as corresponding percentages in the annotated text for each group.

The proportion of total variability due to effect heterogeneity was high (*I²* = 99.5%), which is common in ecological meta-analyses (Senior *et al*., 2016), and indicates that almost all variation stems from true differences in effect sizes rather than sampling variance (Higgins & Thompson, 2002). Within-study variance (effect ID) accounted for 71.7% of the total variability, while between-study variance (article ID) contributed 27.8%, leaving 0.5% attributable to sampling variance.

### 3.2 Univariate moderator analyses

Although the overall responses to both nitrogen enrichment and elevated grazing risk were large and robust, notable underlying heterogeneity in toxin induction existed due to differences between moderator levels (e.g., between different species of *Alexandrium*).

Specifically, *Pseudo-nitzschia* was significantly more induced than *Alexandrium* (Fig. S10). Further, *P. seriata* toxin induction differed from *P. pungens* and all *Alexandrium* species, while several *Alexandrium* species differed from each other (Fig. S11). The induction of different strains were highly variable (Fig. S12), however, most strains appeared in fewer than three cases and could not be statistically resolved. Some of the observed heterogeneity correlated to different culture media used (Fig. S13). However, more than half of all cases used poorly defined “custom” media, making robust inference impossible. Among demand cases that exposed phytoplankton to live grazers, the zooplankton species used also contributed to the underlying heterogeneity (Fig. S14); for example, *Calanus finmarchicus* differed significantly from *Oithona similis* and *Acartia clausi*. This is to be expected as different copepods are known to have different copepodamide profiles and content (Grebner *et al*., 2019; Arnoldt *et al*., 2024). Finally, mean effect size significantly decreased with higher illumination (*p* = 0.004, Table 1:ⅠⅠⅠ, Fig. S15A) and culturing temperature (*p* < 0.0001, Table 1:ⅠⅤ, Fig. S15B).

### 3.3 Interactions between driver and other moderators

In addition to the interaction effect between phytoplankton genus and driver (model ⅠⅠ, Fig. 4), several moderators—including both phytoplankton species and strain, temperature, and illumination level—significantly interacted with driver. However, many of these driver-moderator combination levels contained fewer than three cases (e.g. phytoplankton species and strains) or had large gaps across continuous moderator values (e.g. salinity and temperature), preventing a more detailed analysis. These models are available in subsection 5.2 of the Supplementary code.

### 3.4 Sensitivity analysis

Twelve effect sizes violated Geary’s rule (i.e., had values smaller than 3, the smallest was 1.85) and could potentially bias our overall results. Removing these from the driver (Ⅰ) and driver:phytoplankton genus (ⅠⅠ) models resulted in small overall effect decreases, by 35–36 and 20–85 percentage units respectively (e.g. from 390% to 355% increase), and did not significantly change the results (subsection 6.1 of the Supplementary code).

### 3.5 Publication bias

Visual assessment of the cumulative analysis indicated a potential decline effect (Connell & Leung, 2023) for resource effects over time, decreasing to approximately 8% of the original pooled effect estimated for the first case, while remaining well above a non-significant level (Fig. 5). Demand-driven phycotoxin induction showed a mean effect of approximately 130% increase for over a decade but began to rise from 2015, partly due to the increasing number of experiments on *Pseudo-nitzschia* (Fig. 5). However, there was no evidence of significant time-lag bias or publication bias (p = 0.10 and 0.92, respectively; Table 1:Ⅴ).

**Fig. 5.**
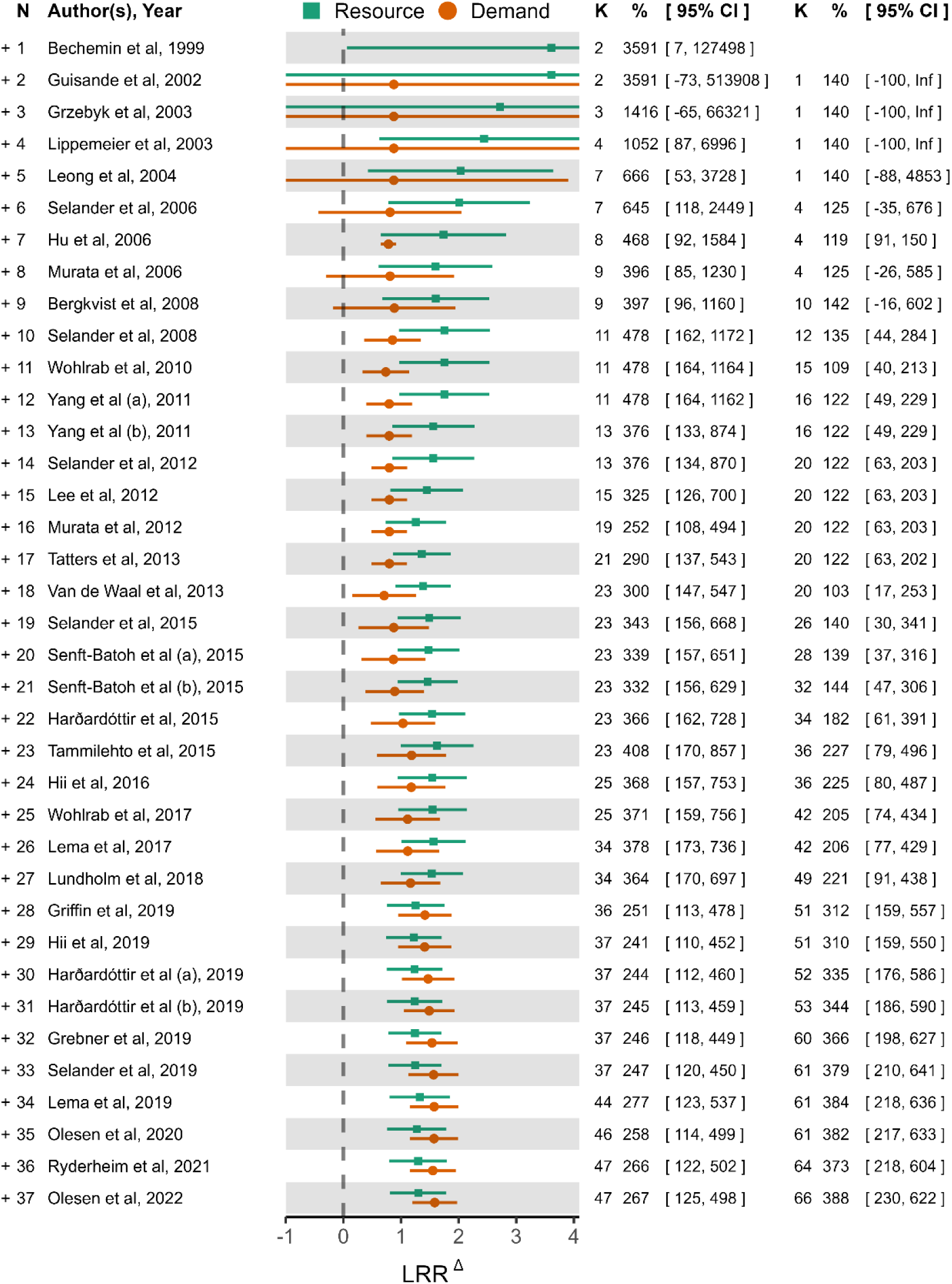
Cumulative forest plot of phycotoxin induction effect size, partitioned by resource and demand effects. The cumulative meta-analysis presents pooled effect sizes as studies are sequentially added in chronological order, illustrating how the overall estimate and its confidence interval (CI) evolve with increasing data inclusion. The filled green squares (resource) and orange circles (demand) represent the pooled effect sizes at each step, with horizontal lines indicating the corresponding 95% CIs. Effect size (x-axis) is the small sample bias-corrected log response ratio (LRR^Δ^) proposed by Lajeunesse (2015). The right-hand table presents estimates as back-transformed percentages, CIs, and cumulative number of effect sizes (k) for clarity. NB: the 95% CIs based on less than approximately 10 effect sizes are not reliable due to the multi-level meta-regression model used; robust estimates were attained once data from Selander et al. (2008) was included, which provided sufficient statistical power for valid hierarchical variance estimation.

## 4. Discussion

We found that nitrogen enrichment (resource) and elevated grazing risk (demand) increased phycotoxin production similarly when pooled across genera. The robustness and consistency of these effects are noteworthy given the considerable underlying heterogeneity in experimental conditions due to phytoplankton species and strains, culturing setups, and experimental conditions used across studies. The response to nitrogen enrichment corroborates a recent meta-analysis on the effects of nutrient limitation on phycotoxins, which demonstrated a similar robust increase (≈170%, 95% CI: 30–470) in paralytic shellfish toxins of *Alexandrium* when grown under phosphorus limitation (Brandenburg *et al*., 2020). Further, our synthesis provides novel insights into the important role of grazers on toxin dynamics in harmful algal bloom (HAB)-forming phytoplankton.

Predation, or simply the risk of predation (Kats & Dill, 1998; Preisser, Bolnick & Benard, 2005; Suraci *et al*., 2016), is a strong selective pressure that has driven the evolution of diverse morphological, chemical, and behavioural defence traits, reducing the likelihood of prey being discovered or consumed. This phenomenon is well-documented across a wide range of organisms (reviewed by, e.g., Abrams, 1990; Kats & Dill, 1998) and can lead to trophic cascade effects at the ecosystem level (Adrian & Schneider-Olt, 1999; Suraci *et al*., 2016; Tiselius & Møller, 2017). Defence adaptations can, in turn, select for counter-adaptations in predators, resulting in an ongoing co-evolutionary arms race between predator and prey (Dawkins & Krebs, 1979; Smetacek, 2001). Planktonic predators exert strong grazing pressure on phytoplankton, which can respond by exhibiting a wide range of inducible defences (Pančić & Kiørboe, 2018). For instance, some species form colonies (Hessen & van Donk, 1993; Lampert, Rothhaupt & von Elert, 1994; Jakobsen & Tang, 2002) or chains (Bergkvist *et al*., 2012; Amato *et al*., 2018) that reduce predator encounter rates or handling efficiency. In marine ecosystems, copepods—the primary mesozooplankton grazers of microalgae—induce various defence traits in their prey. In addition to stimulating toxin production in several phytoplankton taxa, copepods and the chemical cues they exude (copepodamides) can induce bioluminescence in dinoflagellates (Lindström *et al*., 2017; Gonzalo-Valmala et al., in preparation), leading copepods to reject illuminating cells and redirect grazing to less defended prey (Prevett *et al*., 2019). Copepods (and copepodamides) also suppress chain formation in diatoms, which reduces the diatoms’ hydrodynamic signature, decreases their encounter rates with copepods, and consequently reduces grazing risk (Bjærke *et al*., 2015; Rigby & Selander, 2021; Rigby *et al*., 2022). Similarly, motile dinoflagellates, such as *Alexandrium tamarense*, can simultaneously reduce their swimming speed and colony size in response to copepodamides, thereby decreasing their encounter rate with copepods and thus their predation risk (Selander *et al*., 2011). Notably, the intensity of herbivory in pelagic systems is approximately three times more than in terrestrial systems (Cyr & Pace, 1993). Importantly, successful grazing on single-celled algae leads to their immediate and assured death (van Donk, Ianora & Vos, 2011; van Donk *et al*., 2011), unlike most terrestrial plants and some colonial phytoplankton for which partial predation is common. Higher grazing pressure, combined with this binary outcome of grazing, places an even stronger selective pressure on phytoplankton to evolve effective defences against grazers (Dawkins & Krebs, 1979; Harvell, 1990), and highlights the critical role they play in shaping phytoplankton biology.

Manipulating both resource availability and grazing risk in fully factorial designs are rare (e.g., Selander *et al*., 2008; Griffin *et al*., 2019; Ryderheim *et al*., 2021) but particularly effective for teasing apart the relative significance of—and interaction effects between— different drivers. Griffin and colleagues (2019) concluded that grazer exposure was an order of magnitude more effective at inducing *Alexandrium catenella* toxins than any of the nitrogen enrichment treatments, regardless of overall nutrient load. Selander *et al*., (2008) and Ryderheim *et al*., (2021), however, found interactive effects between grazers and nutrient conditions, where toxin production was constrained by nitrogen availability even if the relative increase in toxins caused by grazer exposure were more or less constant and significantly higher than in the absence of grazers. Multifactorial approaches like these are preferable for understanding the complex ecological interactions that facilitate HABs— whether rooted in stoichiometric constraints or evolutionary arms races with predators—and for enabling more effective forecasting and management strategies for these ecologically and economically significant events.

While both *Pseudo-nitzschia* and *Alexandrium* responded similarly in magnitude to nitrogen enrichment, toxin increase in response to grazing risk was ten times higher for *Pseudo-nitzschia* than *Alexandrium*. This pattern is likely due to the lower constitutional domoic acid content in exponentially growing *Pseudo-nitzschia* spp. (Bates, 1998). Thus, even a modest absolute increase yields a large relative effect size. A higher constitutional toxin production rate in dinoflagellates may reflect the higher risk of predation for large, mobile prey such as dinoflagellates compared to that of the small, non-motile *Pseudo-nitzschia* (Selander *et al*., 2011). In addition, diatoms generally have higher growth rates than dinoflagellates (Banse, 1982) and the optimal allocation of resources between growth and defence will likely favor more constitutive defence investments in the more conspicuous dinoflagellates (Feeny 1976). Additionally, while not statistically significant, the mean toxin increase in response to grazing risk was four times greater than that induced by nitrogen enrichment in *Pseudo-nitzschia*.

That relative nitrogen enrichment and elevated grazing risk induced phytotoxins to a similar extent is perhaps surprising given the disparity in published research focus. Nutrient-focused (resource) studies dominate the field (Fig. 6), comprising 76 of the 95 studies fully screened here, compared to only 24 studies on grazing risk (including the five that examined both drivers in both groups). This imbalance likely reflects the strong historical emphasis on marine eutrophication as a driver of HABs (Heisler *et al*., 2008; O’Neil *et al*., 2012) and highlights the need for a more balanced research approach in future studies.

**Fig. 6:**
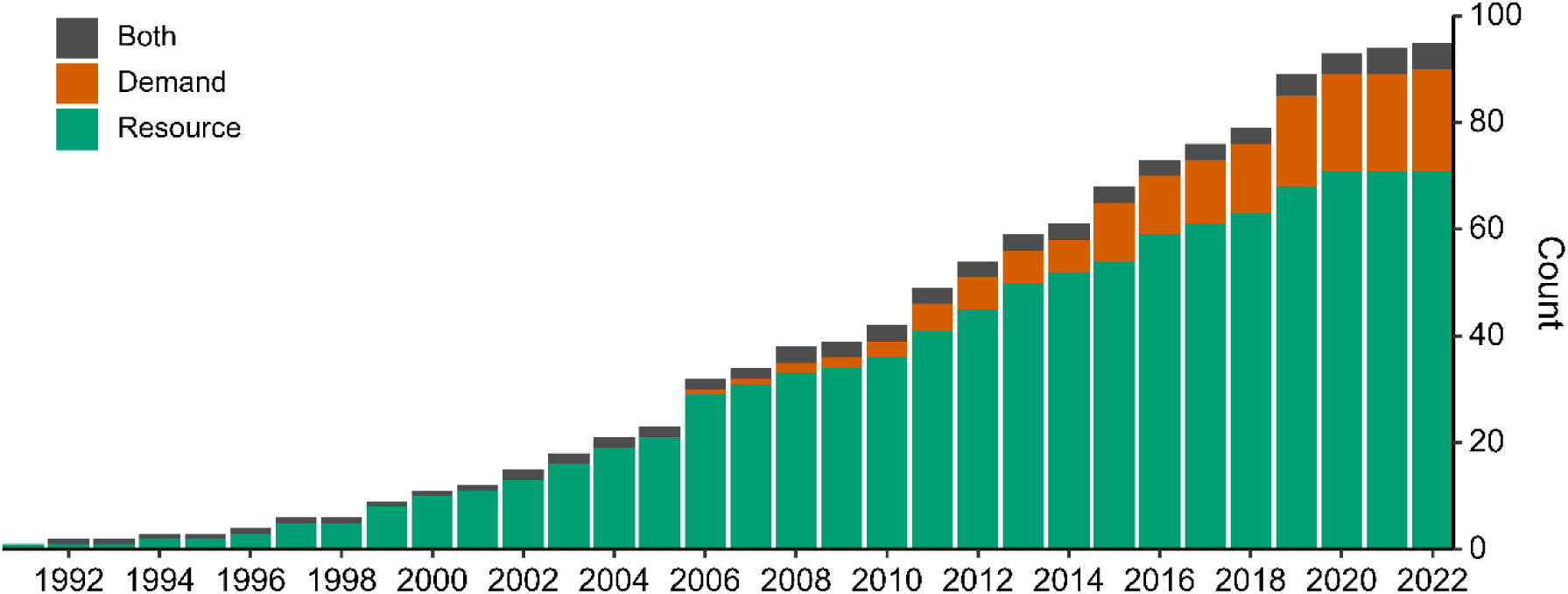
Cumulative number of papers, by publication year (1991-2022) and study type, identified for full-text screening (final screening step of PRISMA-chart, Fig. 2). Papers were categorised as containing resource-(green), demand-(orange), or both resource- and demand-mediated effects (dark grey) on phycotoxins.

### 4.1 Towards a unified theoretical framework

Although terrestrial plant defence hypotheses provide robust frameworks for understanding secondary metabolites (Karban & Baldwin, 1997; Stamp, 2003), these frameworks have seen limited application in explaining toxin production in HAB research. While HAB research has historically emphasised toxin characterisation and distribution patterns in response to resource availability, we argue that drawing insights from the rich theoretical developments in terrestrial plant-herbivore systems offers great potential value. Given our findings, we propose that resource conditions determine the biochemical boundaries for toxin production, as nitrogen-rich phycotoxins require a sufficient supply of nitrogen for synthesis, while grazing risk modulates the extent and timing of toxin production within these constraints to minimise predation risk. This is similar to the hierarchical model of carbon allocation to carbon-rich secondary compounds proposed by Koricheva and colleagues (1998).

Additionally, the life history and grazing vulnerability of prey species likely influences the balance between their constitutive and inducible defence investments. Larger, motile, or slow-growing species that are more apparent to grazers may invest in higher levels of constitutive defences, potentially complemented by inducible defences (Feeny, 1976). In contrast, smaller or less conspicuous species may primarily rely on inducible defences that are up-regulated only under elevated grazing risk. Moreover, HAB-forming species serve as powerful model systems for studying plant-grazer dynamics, thanks to their short generation times, ease of laboratory cultivation, and suitability for multifactorial experimental designs that are often impractical in terrestrial plant models. Moving forward, HAB research should include experimental designs and interpretative frameworks that explicitly address both resource-driven and grazer-driven responses. While distinct from terrestrial systems, marine phytoplankton toxin dynamics can be better understood through careful consideration of resource constraints, the ecology of the species, and grazing pressure.

### 4.2 Limitations

Our analysis was confined to two genera of marine HAB species, precluding direct generalisation to limnic systems and in particular to cyanobacteria which are not represented in the current study. A comprehensive synthesis across all marine HAB-forming genera would strengthen our conclusions, however, insufficient primary research on several taxonomic groups currently prevents such analysis. We were unable to include growth rate as a moderator variable in our analysis, partly due to it being poorly reported in primary studies, despite growing evidence of its significance in modulating cell specific toxin content (Anderson *et al*., 1990; John & Flynn, 2000; Guisande *et al*., 2002; Pourdanandeh *et al*., 2025). Growth dynamics may be less relevant when the primary interest is volumetric toxin concentration, a metric of direct relevance to stakeholders such as shellfish farmers.

However, we suggest that future studies of mechanisms that influence toxin production *per se* (as opposed to cell specific toxin content) should include growth rate as a covariate in formal analyses, or use metrics such as the cell specific toxin production rate (Anderson *et al*., 1990) as response variable. Finally, although we used one of the few statistically sound methods for detecting publication bias in datasets characterized by high heterogeneity and non-independence (Nakagawa *et al*., 2022), its results should be interpreted with caution.

Simulation studies indicate that while most analytical methods maintain low Type I error rates, this often comes at the cost of increased Type II error rates, potentially leading to conservative underestimations of publication bias (Fernández-Castilla *et al*., 2021; Rodgers & Pustejovsky, 2021).

## 5. Conclusions

1. This synthesis reflects the ongoing shift from viewing harmful algal toxin production primarily through the lens of nutrient-mediated resource dynamics to recognising the strong influence of grazers as selective agents of plant defences.
2. We show that grazers rival the aggregated phycotoxin induction potential of nitrogen enrichment and supersede it for the amnesic shellfish toxin-producing diatom *Pseudo-nitzschia*. Toxin production appears to be under tight control by the level of threat in both genera, but more so in *Pseudo-nitzschia*.
3. Grazer-induced toxin production may be contributing to the enigmatic fluctuations of toxins observed in situ, which hamper the development of HAB forecasting models.
4. Our findings motivate a more balanced effort in future HAB research, integrating resource-based and demand-based factors while linking experimental observations to the well-established theoretical frameworks of plant defence.
5. By integrating both perspectives and designing experiments to test predictions derived from plant defence models, we can develop a better understanding of the drivers and mechanisms behind toxic harmful algal blooms, ultimately improving our ability to predict and manage these ecologically and economically impactful events.

## Acknowledgements

This work was funded by the Swedish Research Council grant to **E.S** (VR 2019-05238). We thank Jonathan N. Havenhand and Lars Gamfeldt for their valuable comments and feedback, which greatly improved this manuscript.

---

**Fig. S1:**
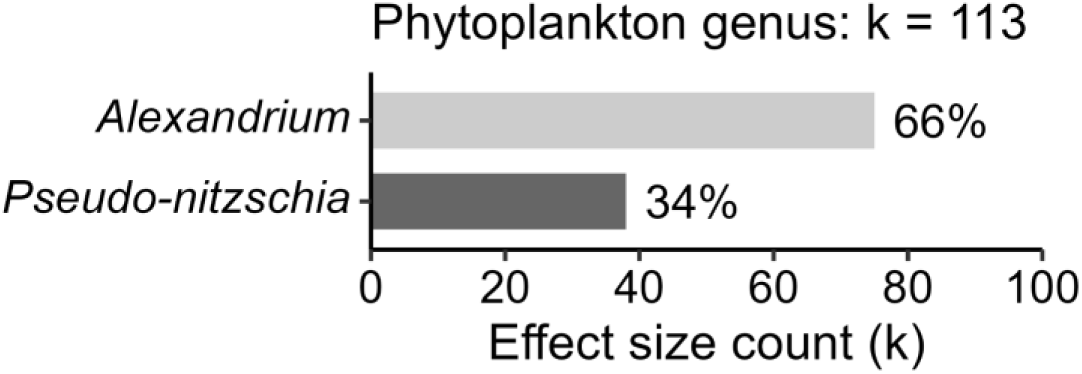
Effect sizes (k) distributed between the two phytoplankton genera *Alexandrium* and *Pseudo-nitzschia*.

**Fig. S2:**
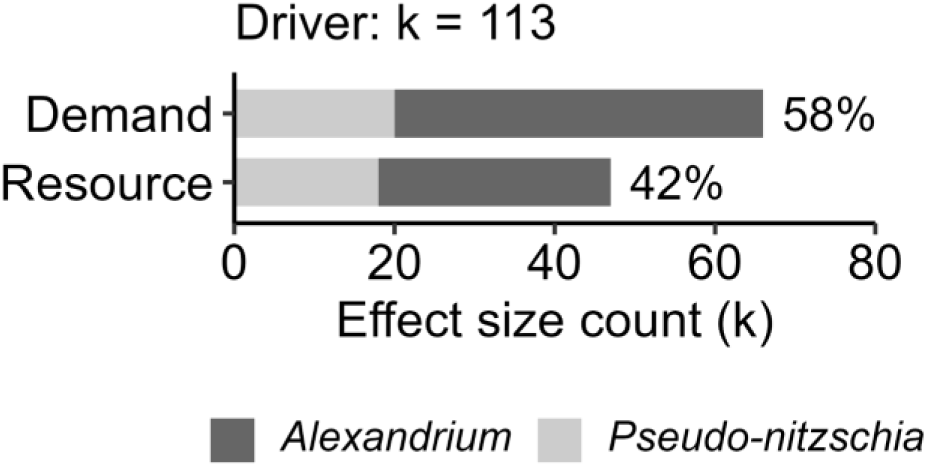
Effect sizes (k) distributed between levels of moderator driver (demand & resource), coloured by phytoplankton genus (*Alexandrium* & *Pseudo-nitzschia*).

**Fig. S3:**
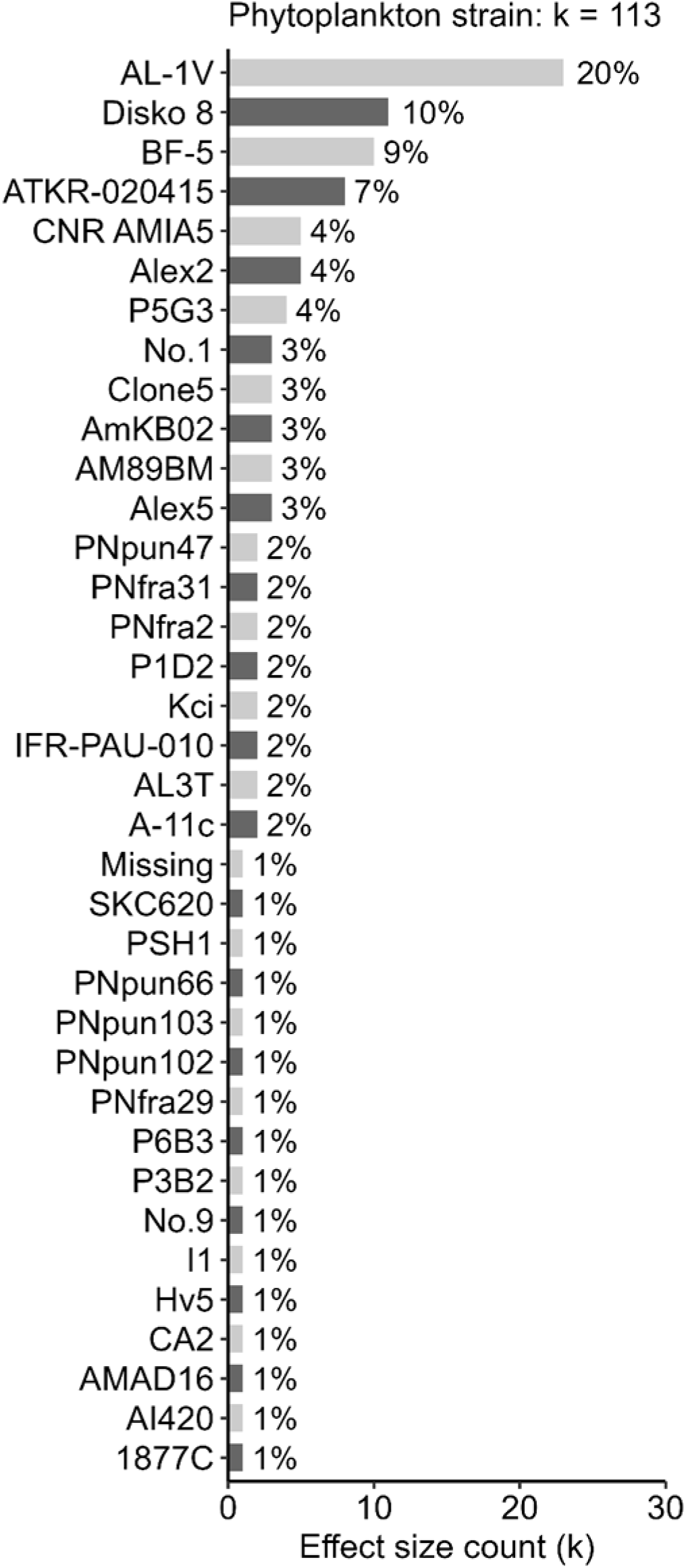
Effect sizes (k) distributed among phytoplankton strains.

**Fig. S4:**
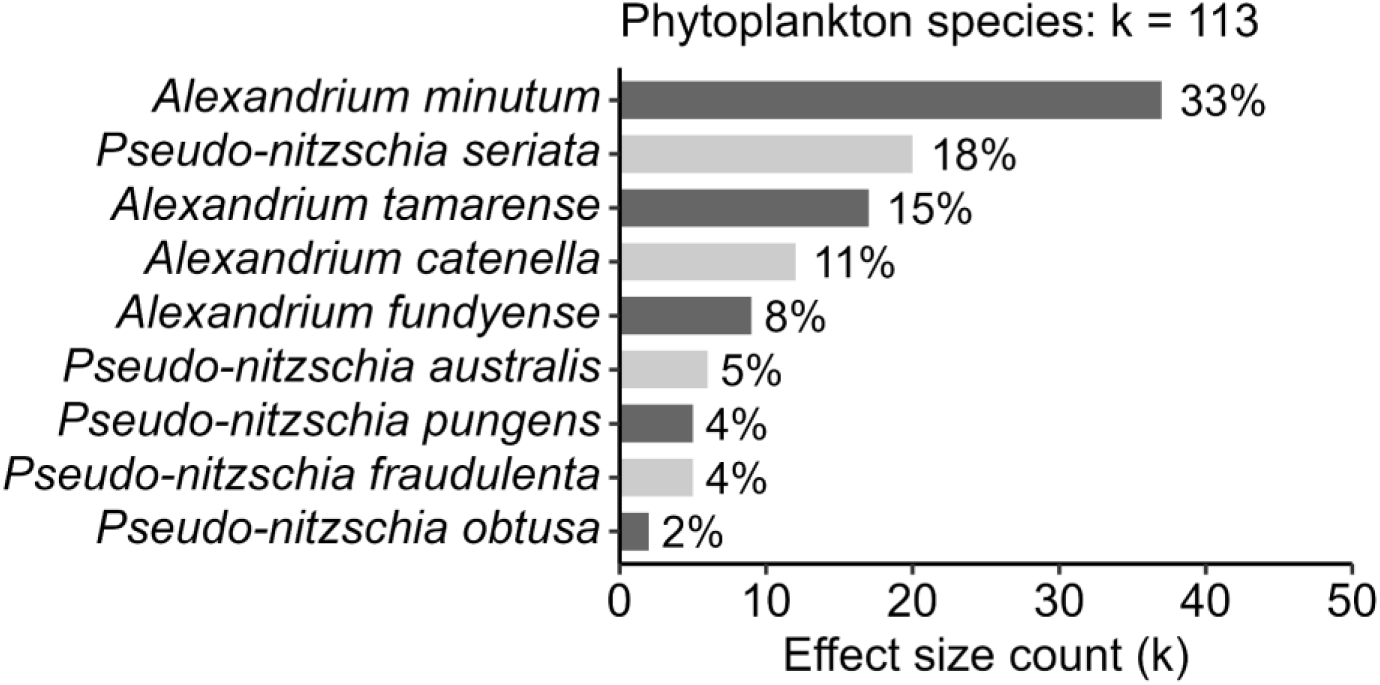
Effect sizes (k) distributed among phytoplankton species.

**Fig. S5:**
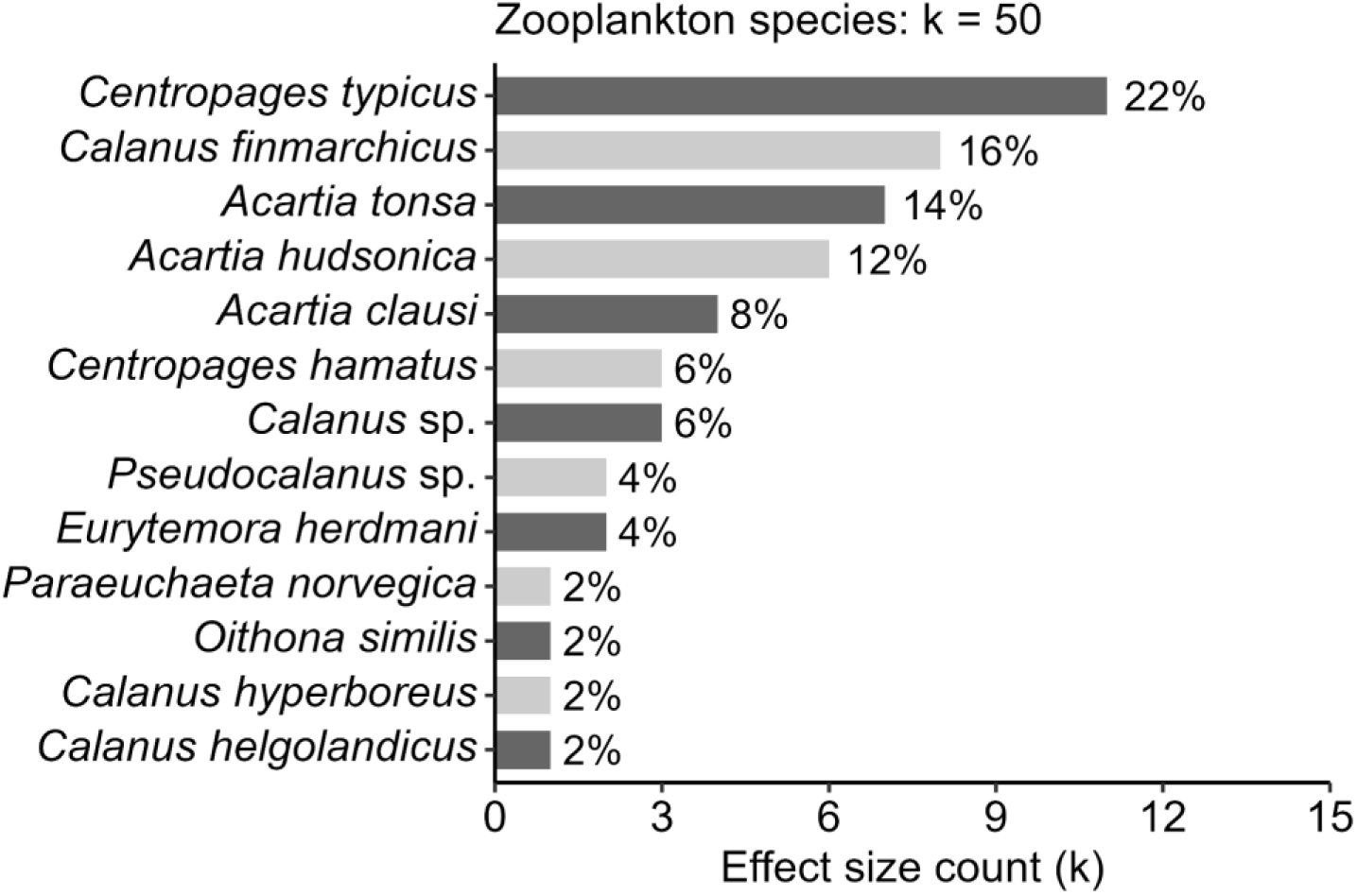
Effect sizes (k) from studies that exposed phytoplankton to live zooplankton, distributed among zooplankton (copepod) species.

**Fig. S6:**
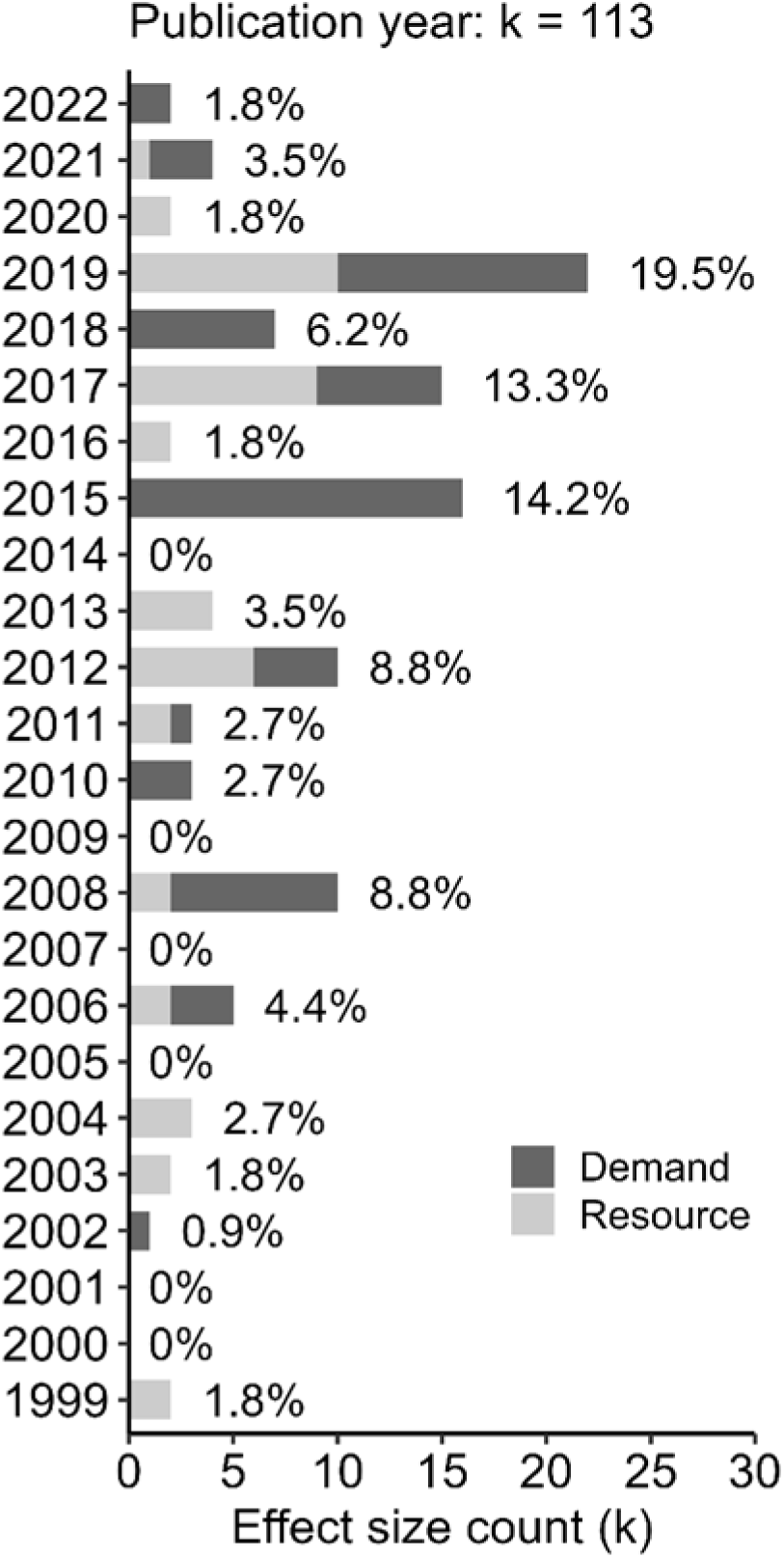
Effect sizes (k) distributed over time, coloured by experiment type/driver (demand & resource).

**Fig. S7:**
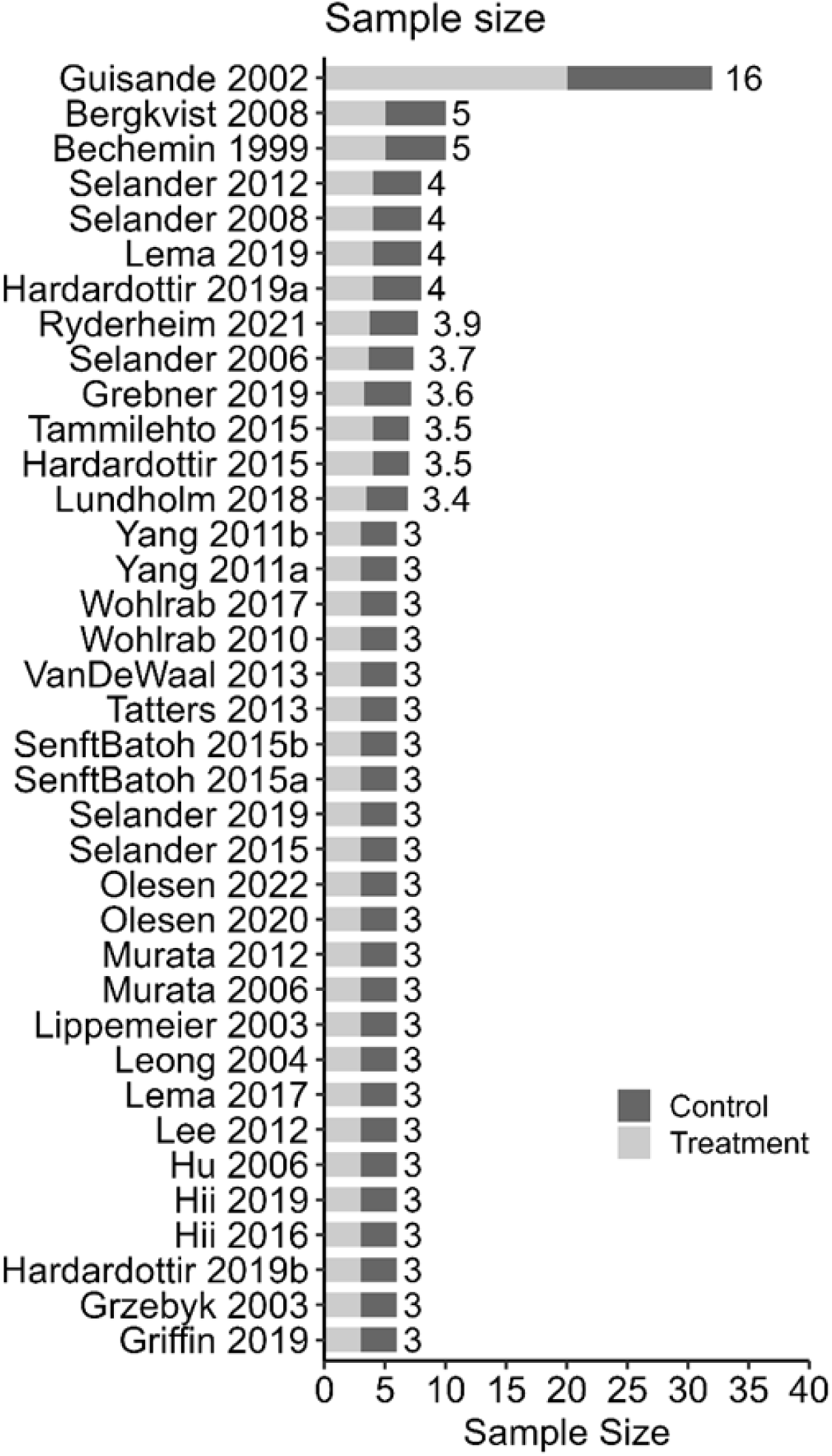
Total sample size of studies included in the analysis, coloured by experimental group (control & treatment). For experimental contrasts between different nutrient levels (yielding different N:P ratios) of resource papers/effects, the treatment with the lower N:P ratio was consistently defined as the control. The numbers next to each bar denotes the average sample size for controls and treatments in each study.

**Fig. S8:**
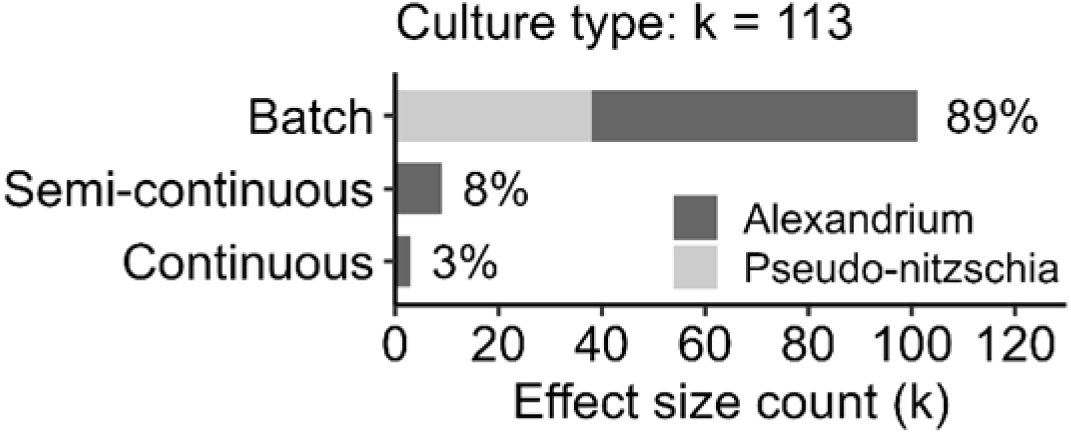
Effect sizes (k) distributed between levels of moderator culture type, partitioned by phytoplankton genus (Alexandrium & Pseudo-nitzschia).

**Fig. S9:**
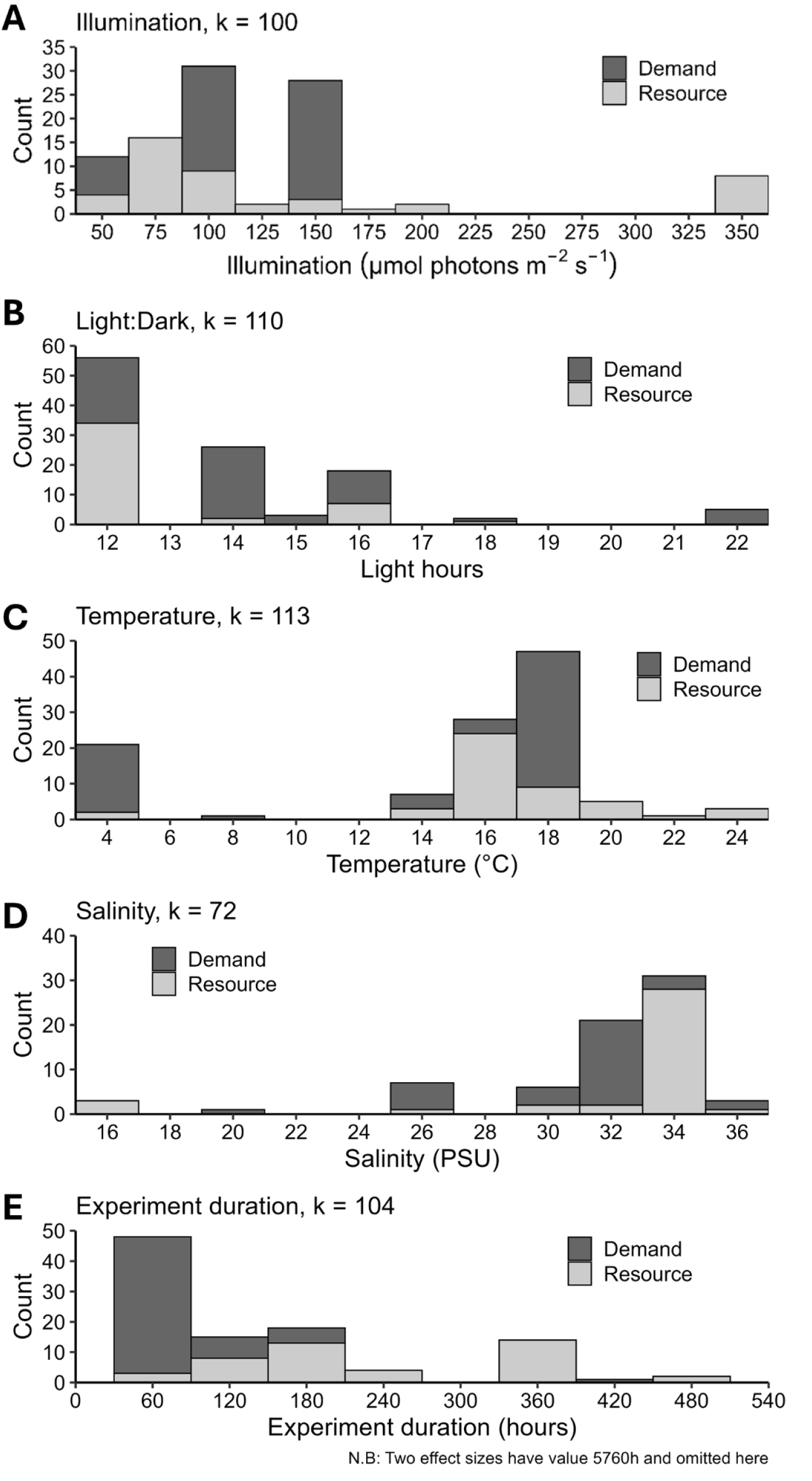
Stacked histogram of effect sizes (k) distributed over continuous moderators **(A)** light intensity, **(B)** light:dark cycle, **(C)** temperature, **(D)** salinity, **(E)** and duration of experiment. Bars are coloured by experiment type/driver (demand & resource). Note that two cases were omitted from **(E)** as they ran for more than 10x longer (5760h) than the second longest experiments (480h).

**Fig. S10:**
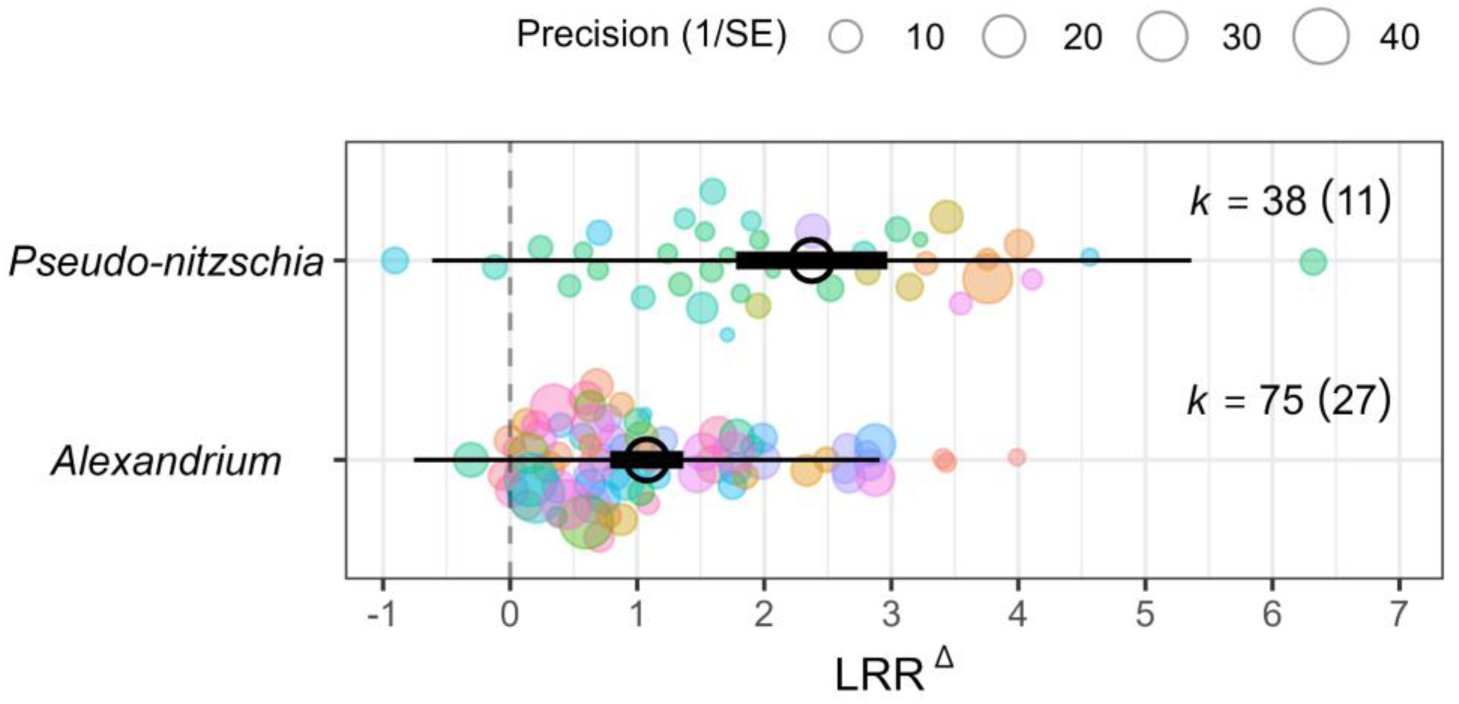
Effects of relative nitrogen enrichment (resource) or elevated grazing risk (demand) on phycotoxin induction (LRR^Δ^), separated by phytoplankton genus (*Alexandrium* and *Pseudo-nitzschia*). LRR^Δ^ is the small sample bias-corrected log response ratio proposed by Lajeunesse (2015). Empty black circles represent mean effect sizes, thick black lines denote 95% confidence intervals (95% CI) for the mean effects, and thin black lines indicate 95% prediction intervals (95% PI; where 95% of new effect sizes are expected to fall with repeated sampling of the literature). Coloured circles denote individual effect sizes (k) from a given number of studies (n), sized inversely proportional to their sampling error (1/SE), and coloured by their study ID.

**Fig. S11:**
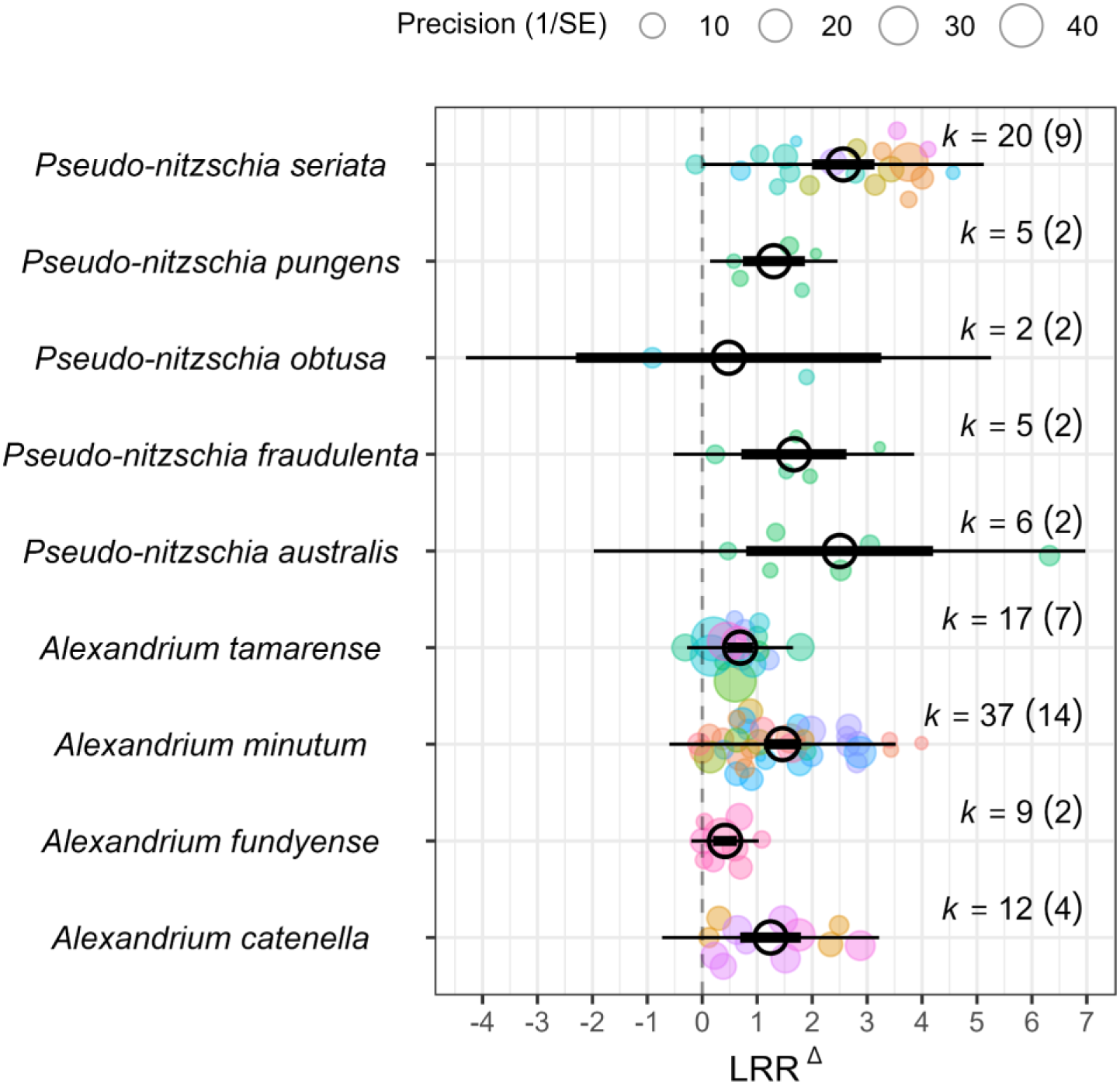
Effects of relative nitrogen enrichment (resource) or elevated grazing risk (demand) on phycotoxin induction (LRR^Δ^), separated by phytoplankton species within genera *Alexandrium* and *Pseudo-nitzschia*. LRR^Δ^ is the small sample bias-corrected log response ratio proposed by Lajeunesse (2015). Empty black circles represent mean effect sizes, thick black lines denote 95% confidence intervals (95% CI) for the mean effects, and thin black lines indicate 95% prediction intervals (95% PI; where 95% of new effect sizes are expected to fall with repeated sampling of the literature). Coloured circles denote individual effect sizes (k) from a given number of studies (n), sized inversely proportional to their sampling error (1/SE), and coloured by their study ID.

**Fig. S12:**
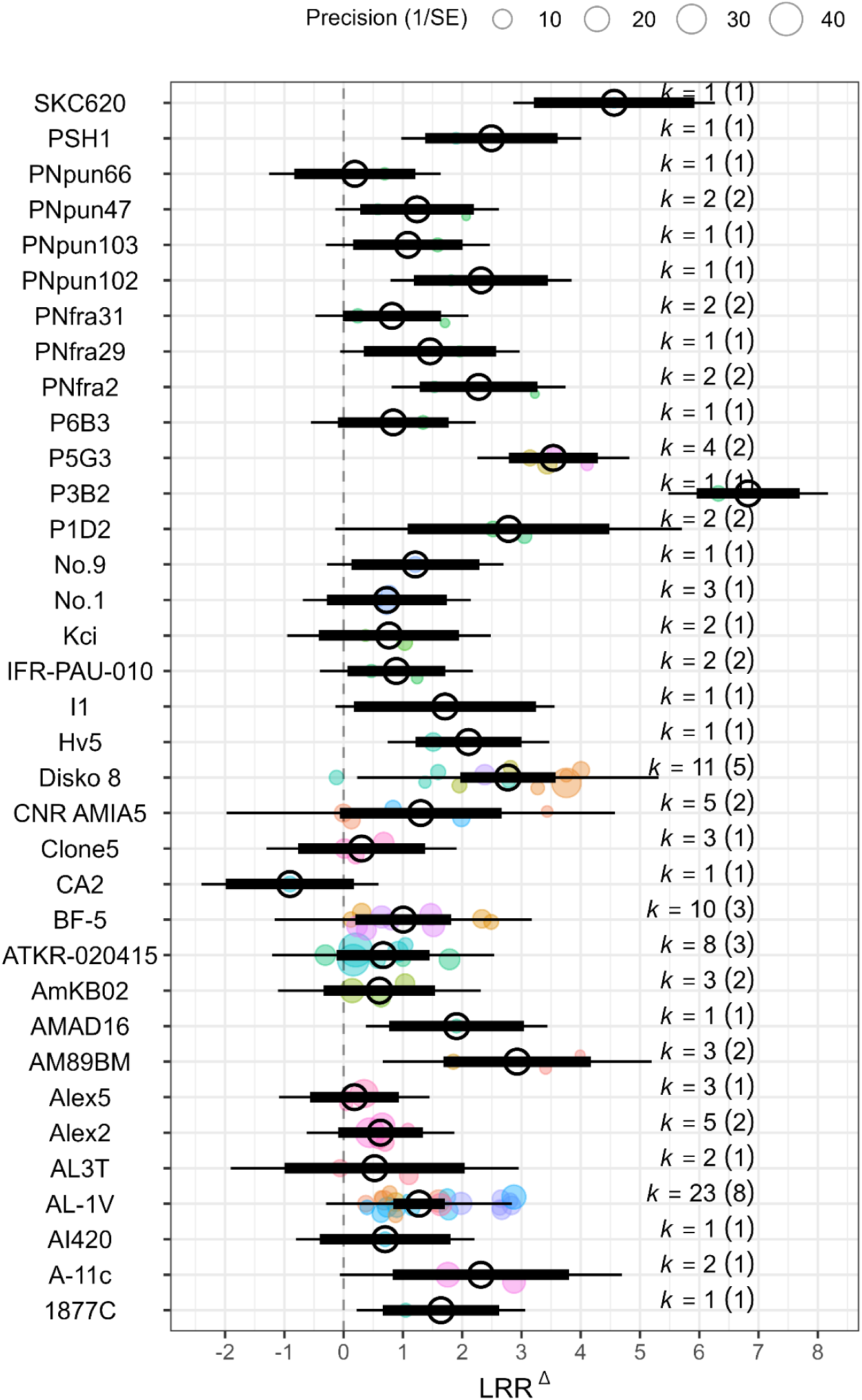
Effects of relative nitrogen enrichment (resource) or elevated grazing risk (demand) on phycotoxin induction (LRR^Δ^), partitioned among phytoplankton strains. LRR^Δ^ is the small sample bias-corrected log response ratio proposed by Lajeunesse (2015). Empty black circles represent mean effect sizes, thick black lines denote 95% confidence intervals (95% CI) for the mean effects, and thin black lines indicate 95% prediction intervals (95% PI; where 95% of new effect sizes are expected to fall with repeated sampling of the literature). Coloured circles denote individual effect sizes (k) from a given number of studies (n), sized inversely proportional to their sampling error (1/SE), and coloured by their study ID. separated by culture medium used.

**Fig. S13:**
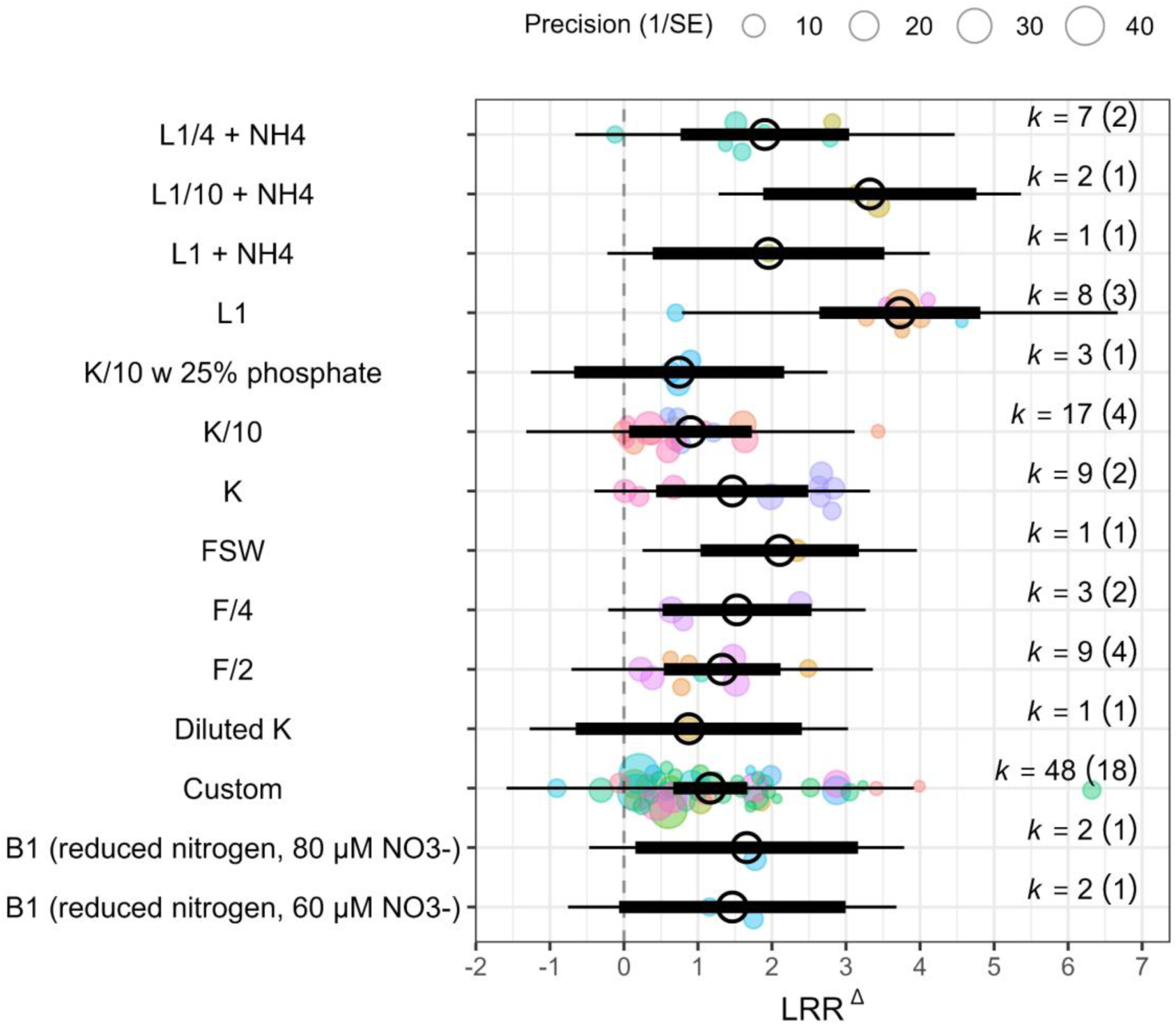
Effects of relative nitrogen enrichment (resource) or elevated grazing risk (demand) on phycotoxin induction (LRR^Δ^), separated by culture medium used. LRR^Δ^ is the small sample bias-corrected log response ratio proposed by Lajeunesse (2015). Empty black circles represent mean effect sizes, thick black lines denote 95% confidence intervals (95% CI) for the mean effects, and thin black lines indicate 95% prediction intervals (95% PI; where 95% of new effect sizes are expected to fall with repeated sampling of the literature). Coloured circles denote individual effect sizes (k) from a given number of studies (n), sized inversely proportional to their sampling error (1/SE), and coloured by their study ID.

**Fig. S14:**
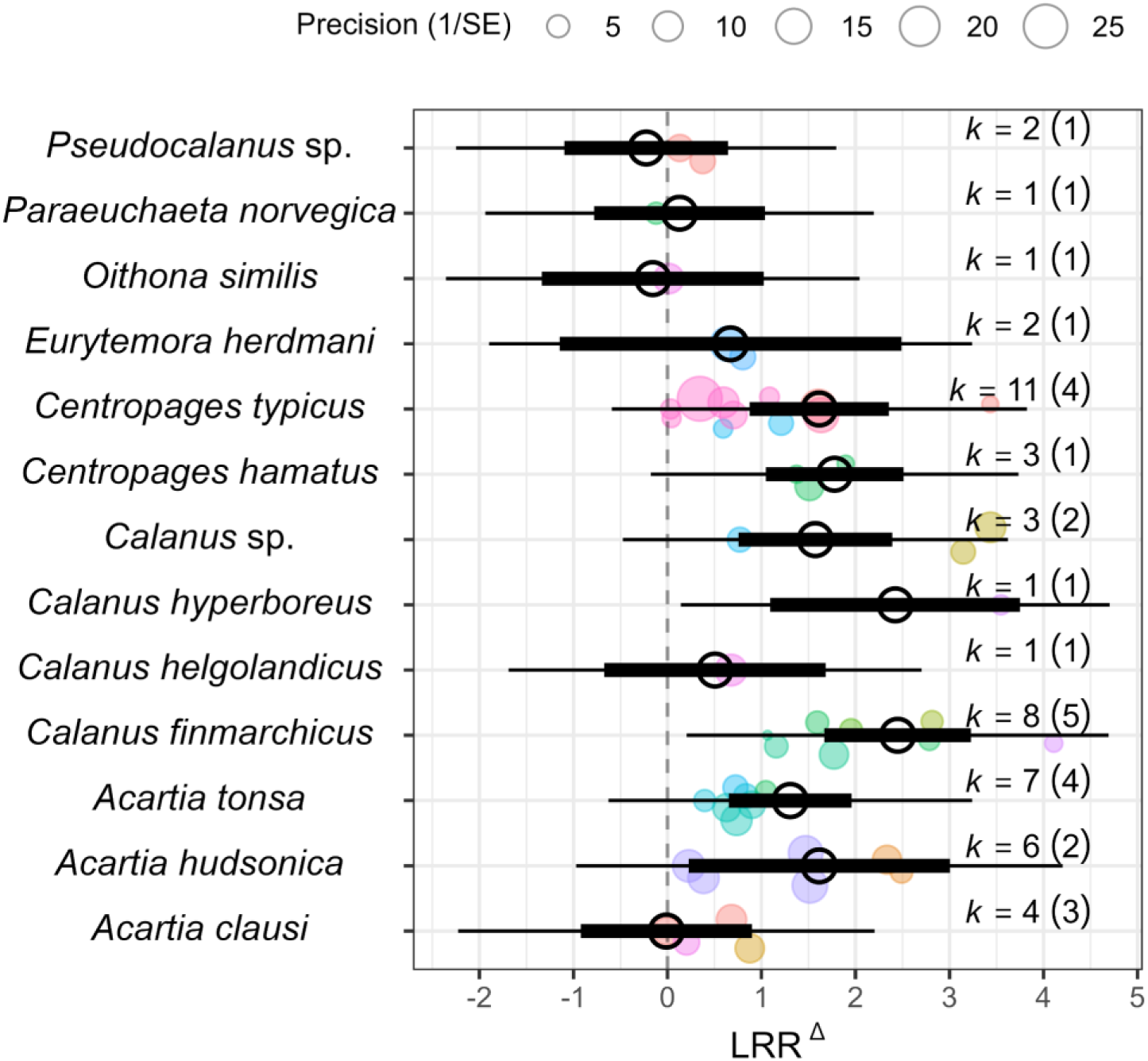
Effects of elevated grazing risk (demand) on phycotoxin induction (LRR^Δ^), separated by the copepod species used. LRR^Δ^ is the small sample bias-corrected log response ratio proposed by Lajeunesse (2015). Empty black circles represent mean effect sizes, thick black lines denote 95% confidence intervals (95% CI) for the mean effects, and thin black lines indicate 95% prediction intervals (95% PI; where 95% of new effect sizes are expected to fall with repeated sampling of the literature). Coloured circles denote individual effect sizes (k) from a given number of studies (n), sized inversely proportional to their sampling error (1/SE), and coloured by their study ID.

**Fig. S15:**
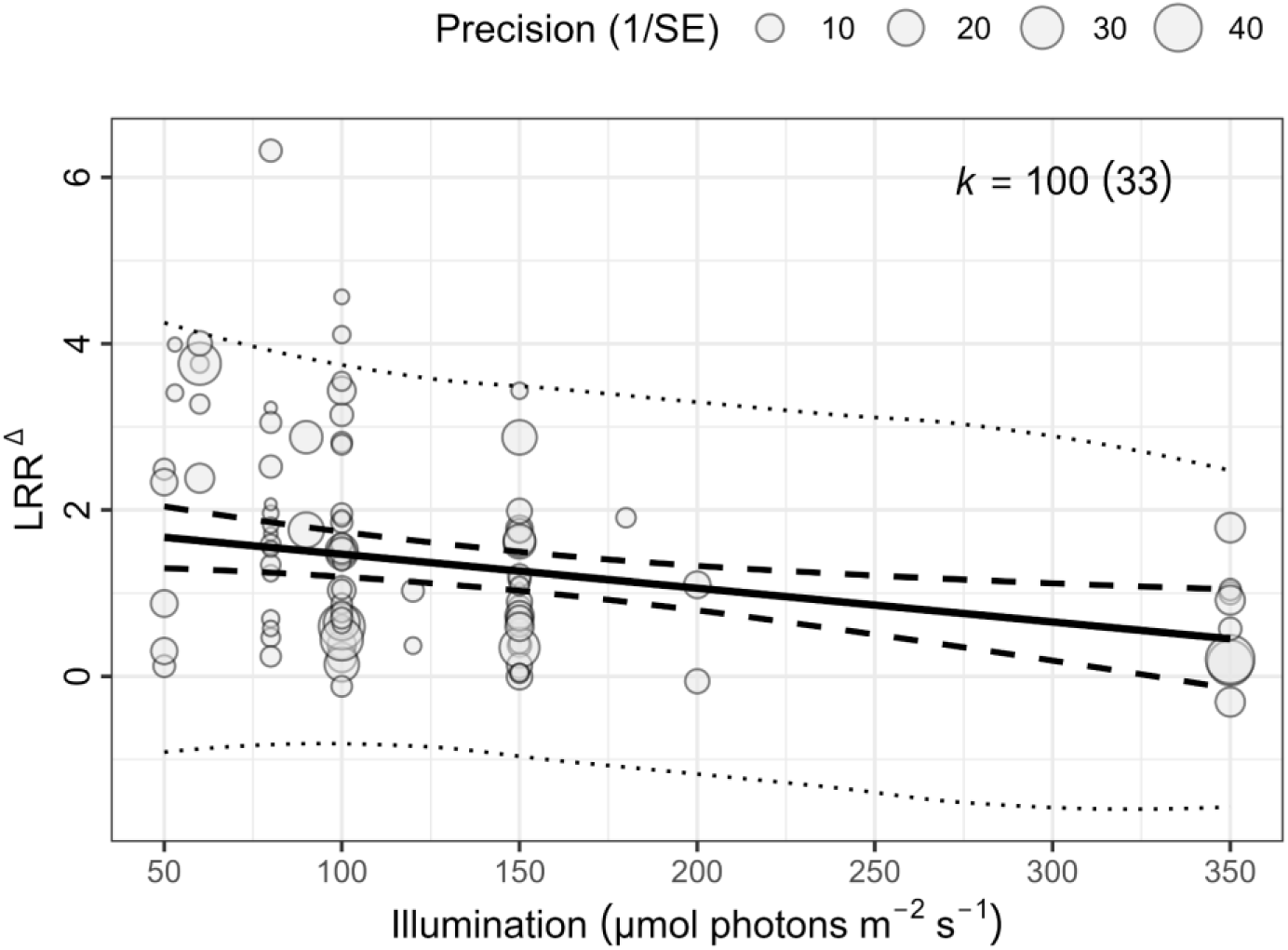
Effects of relative nitrogen enrichment (resource) or elevated grazing risk (demand) on phycotoxin induction (LRR^Δ^), as a function of experimental light intensity. LRR^Δ^ is the small sample bias-corrected log response ratio proposed by Lajeunesse (2015). The solid black line is the fitted linear relationship from meta-regression, thick dashed lines denote 95% confidence interval (95% CI) for the fit, and the thin dotted lines indicate 95% prediction interval (95% PI; where 95% of new effect sizes are expected to fall with repeated sampling of the literature). Circles denote individual effect sizes (k) from a given number of studies (n), sized inversely proportional to their sampling error (1/SE).

